# Dopamine neuron dysfunction and loss in the *Prkn*R275W mouse model of Juvenile Parkinsonism

**DOI:** 10.1101/2023.09.26.559326

**Authors:** Maria Regoni, Letizia Zanetti, Martina Sevegnani, Chiara Domenicale, Stefano Magnabosco, Jyoti C. Patel, Megan K. Fernandes, Elena Monzani, Stefano Comai, Cherchi Laura, Andrea Ciammola, Flavia Valtorta, Michele Morari, Giovanni Piccoli, Margaret E. Rice, Jenny Sassone

**Affiliations:** San Raffaele Scientific Institute, Milan, Italy; Vita-Salute San Raffaele University, Milan, Italy; Dulbecco Telethon Institute, Laboratory of Biology of Synapse, Center for Integrative Biology (CIBIO), University of Trento, Trento, Italy; Department of Neuroscience and Rehabilitation, University of Ferrara, Ferrara, Italy; Department of Neurosurgery, New York University Grossman School of Medicine, New York, NY, USA; Department of Biomedical Sciences, University of Padova, Padova, Italy; IRCCS Istituto Auxologico Italiano, Department of Neurology and Laboratory of Neuroscience, Milan

**Keywords:** PARKIN, neurodegeneration, mouse model

## Abstract

Mutations in the *PRKN* gene encoding the protein PARKIN cause Autosomal Recessive Juvenile Parkinsonism (ARJP). Harnessing this mutation to create an early-onset Parkinson’s disease (PD) mouse model would provide a unique opportunity to clarify the mechanisms involved in the neurodegenerative process and lay the groundwork for the development of neuroprotective strategies. We created a knock-in mouse carrying the homozygous *Prkn*R275W mutation, which is the missense mutation with the highest allelic frequency in *PRKN* patients. In *Prkn*R275W mice, we analysed the anatomical and functional integrity of the nigrostriatal pathway, including striatal DA content and evoked striatal dopamine (DA) release, as well as the motor phenotype.

We report here that *Prkn*R275W mice show early DA neuron dysfunction, age-dependent loss of DA neurons in the substantia nigra, decreased DA content and stimulus-evoked DA release in the striatum, and progressive motor impairment. Together, these data show that the *Prkn*R275W mouse recapitulates key features of ARJP. Thus, these studies fill a critical need in the field by introducing a promising new PD model in which to study causative mechanisms of the disease, as well as test therapeutic strategies.

## Introduction

Mutations in the *PRKN* gene, which encodes for the protein PARKIN, are the most common cause of Autosomal Recessive Juvenile Parkinsonism (ARJP) (*PRKN*, OMIM 600116).^1^ The disease is characterized by the progressive loss of dopamine (DA) neurons in the substantia nigra pars compacta (SNc),^2^ which leads to DA depletion in the striatum and appearance of typical motor features such as bradykinesia, resting tremor and rigidity.^3,4^ Median age at onset is ∼31 years.^3,4^ More than 100 different *PRKN* mutations have been found, spread along all 12 exons of the gene. The most frequent mutations are exon rearrangements resulting in premature stop codons, degradation of the transcript and absence of PARKIN protein. Disease-causing mutations also include single-nucleotide substitutions resulting in missense, nonsense, or splice-site mutations.^5^ (https://www.mdsgene.org).

In initial attempts to create mechanistic models of the disease, germline *Prkn* exon deletions in mice were introduced. Five different *Prkn* knockout (KO) mice were generated;^6^ these mice show mitochondrial dysfunction and broad spectrum proteomic changes. However, neither overt DA neuron loss nor a parkinsonian phenotype was observed in these models.^7^ The reason why *Prkn* KO models do not recapitulate the phenotype expected from similar mutations in the human orthologue is unknown. This is unlikely to be due to the shorter lifespan and different biology of the mouse compared to human, since *Prkn* deletion in adult mice via lentiviral delivery of Cre-recombinase in midbrain leads to progressive DA neuron loss.^8^ Moreover, selective expression of the human variant PARKINQ311X in DA neurons leads to age-dependent DA neuron degeneration in the SNc.^9,10^ It is therefore possible that *Prkn* deletion from birth can lead to genetic compensation, a phenomenon known to frequently occur in response to gene KO at germline levels.^11–13^

To genetically reproduce the human ARJP condition in mice without changing DNA copy number and avoiding genetic compensation, we created a knock-in mouse model carrying the p.Arg275Trp (R275W) mutation, i.e. the missense mutation with the highest allelic frequency in *PRKN* patients.^5^ In the *Prkn* mouse orthologue, the amino acid R274 corresponds to human R275, thus we inserted the R274W mutation in the *Prkn* gene. The mutation introduced in the murine DNA allows a physiological PARKIN expression under the endogenous promoter and genetically reproduces the expression of missense PARKIN variants in the ARJP patients. In this study, we analysed the anatomical and functional integrity of the nigrostriatal pathway of homozygous *Prkn*R275W mice including striatal DA content and evoked striatal DA release, as well as their motor phenotype.

## Materials and methods

The full methodology is provided in the Supplementary material.

## Data availability

All data generated in this study are available upon reasonable request.

## Results

### Generation of the *Prkn*R275W mouse

Using the CRISPR/Cas9 Technology, we created a new knock-in mouse model expressing the *PRKN* mutation p.Arg275Trp (R275W), that is the missense mutation with the highest allelic frequency in ARJP patients (https://www.mdsgene.org/d/1/g/4). The R275 amino acid in the human cDNA corresponds to the R274 amino acid in the *Prkn* mouse orthologue (Ensembl gene ID: ENSMUSG00000023826). Alongside the exchange of the first nucleotide in the R274 codon of the *Prkn* gene, two silent mutations were introduced close to the site of the mutation to create a novel AflIII restriction site that allows for genotyping of the mutated allele (**Fig. 1A-C**).

**Figure 1.**
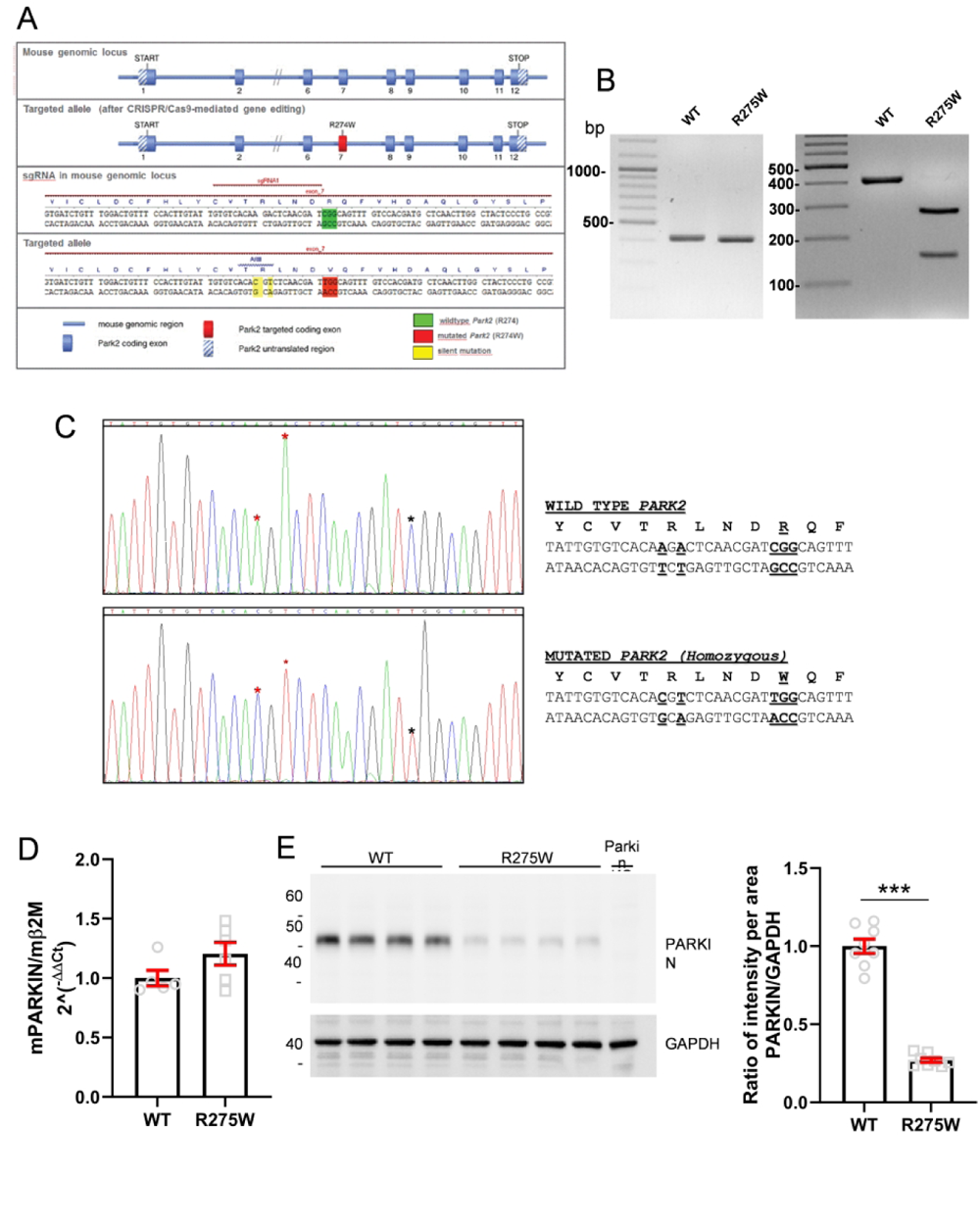
Generation of the *Prkn*R275W mouse. **(A)** The image shows the strategy used for generating a constitutive knock-in of a point mutation in the *Prkn* gene using the CRISPR/Cas9 technology. In the *Prkn* mouse orthologue (Ensembl gene ID: ENSMUSG00000023826), we introduced the R274W missense mutation. In the *Prkn* mouse orthologue the amino acid R274 corresponds to human R275. Alongside the exchange of the first nucleotide on the R274 codon, two silent mutations were introduced close to the site of the mutation to create a novel AflIII restriction site that allows for genotyping of the mutated allele. **(B)** Left: representative image of the result of a PCR with primers that amplify a region the exon 7 in the *Prkn* locus containing the mutated codon. The expected fragment was 424bp. Right: AflIII digestion of this PCR product resulted in cleavage of the 424bp PCR product in two fragments (279bp and 145bp) selectively in the mutated allele. DNA ladder is 100bp. **(C)** Sanger sequence analysis of exon 7 in the *Prkn* locus in genomic DNA from WT mouse and mouse homozygous for the R275W mutations. The black asterisk labels the missense mutation. Red asterisks indicate the silent mutation introduced to create the AflIII restriction site. **(D)** Quantitative real-time PCR performed on mRNA prepared from total mouse brains at 1 month of age with TaqMan probes specific for *Prkn* mRNA and for the housekeeping *Beta-2 microglobulin* (*β2M*) mRNA. The level of endogenous *Prkn* mRNA was normalized on *β2M*. The levels of *Prkn* mRNA in WT and homozygous *Prkn*R275W mice were virtually identical (n = 5 brains from WT mice, n = 6 brains from *Prkn*R275W mice, two-tailed unpaired t-test; p > 0.05). **(E)** Representative western blot performed on lysates from mouse brains at 1 month of age with an antibody specific for PARKIN (monoclonal antibody P6248 - Sigma-Aldrich). *Prkn*R275W mice showed significantly reduced endogenous PARKIN protein levels when compared with WT (1 mo WT 1.00 ± 0.05 *vs.* 1 mo *Prkn*R275W 0.27 ± 0.01, two-tailed unpaired t-test; ***p < 0.001; t = 15.22, df = 14, n= 8 brains from WT mice, n = 8 brains from *Prkn*R275W mice). Negative control is a brain protein lysate from *Prkn* KO mouse.

Homozygous R275W mice were born at expected Mendelian ratios, they appeared normal and were viable and fertile. To assess the expression of the edited *Prkn* gene we performed a real-time PCR on total brain lysates at 1 month of age. *Prkn* mRNA levels were similar between wild-type (WT) and R275W mice, indicating that the missense mutation *Prkn*R275W did not induce *Prkn* mRNA downregulation (**Fig. 1D**). We then determined the levels of PARKIN protein in total brain lysates. R275W mice showed significantly reduced levels of endogenous PARKIN protein when compared with WT mice (**Fig. 1E** and **Fig. S1**-**2**). This is in agreement with previous studies showing that R275W mutation in PARKIN protein results in a conformational alteration^14^ and loss-of-function phenotype.^15^

### Dysfunction of SNc DA neurons in young (1-month-old) *Prkn*R275W mice

To elucidate the impact of R275W mutation on DA neurons, we counted the number of tyrosine hydroxylase positive (TH+) neurons and total neurons (identified by Nissl staining, Nissl+) in the SNc of WT and *Prkn*R275W mice using unbiased stereology.

At 1 month of age, WT and *Prkn*R275W mice had similar TH+ and Nissl+ neuron counts (**Fig. 2A**), indicating the absence of overt nigrostriatal degeneration. However, examination of SNc DA neurons from *Prkn*R275W mice at higher magnification revealed cytoplasmic vacuolizations which were not present in WT mice. The average number of vacuoles per cell was 0.4 (range 0–2) in WT SNc DA neurons and 2.8 (range 0–10) in *Prkn*R275W SNc DA neurons (**Fig. 2B**). As a negative control, we also analyzed DA neurons in the ventral tegmental area (VTA), a brain area that does not undergo extensive degeneration in PD. One to two vacuoles per cell were observed in a small subset of DA neurons in the VTA from WT and *Prkn*R275W mice at 1 month of age, with no difference between genotypes (**Fig. S3**). Therefore, cytoplasmic vacuolization is a feature of the SNc DA neurons of *Prkn*R275W mice; similar cytoplasmic vacuolation has been found in isogenic human DA neurons bearing PARKIN mutations^16^ and in SNc DA neurons of the PARKINQ311X mouse.^10^ This result suggests early dysfunction of SNc DA neurons in *Prkn*R275W mice.

**Figure 2.**
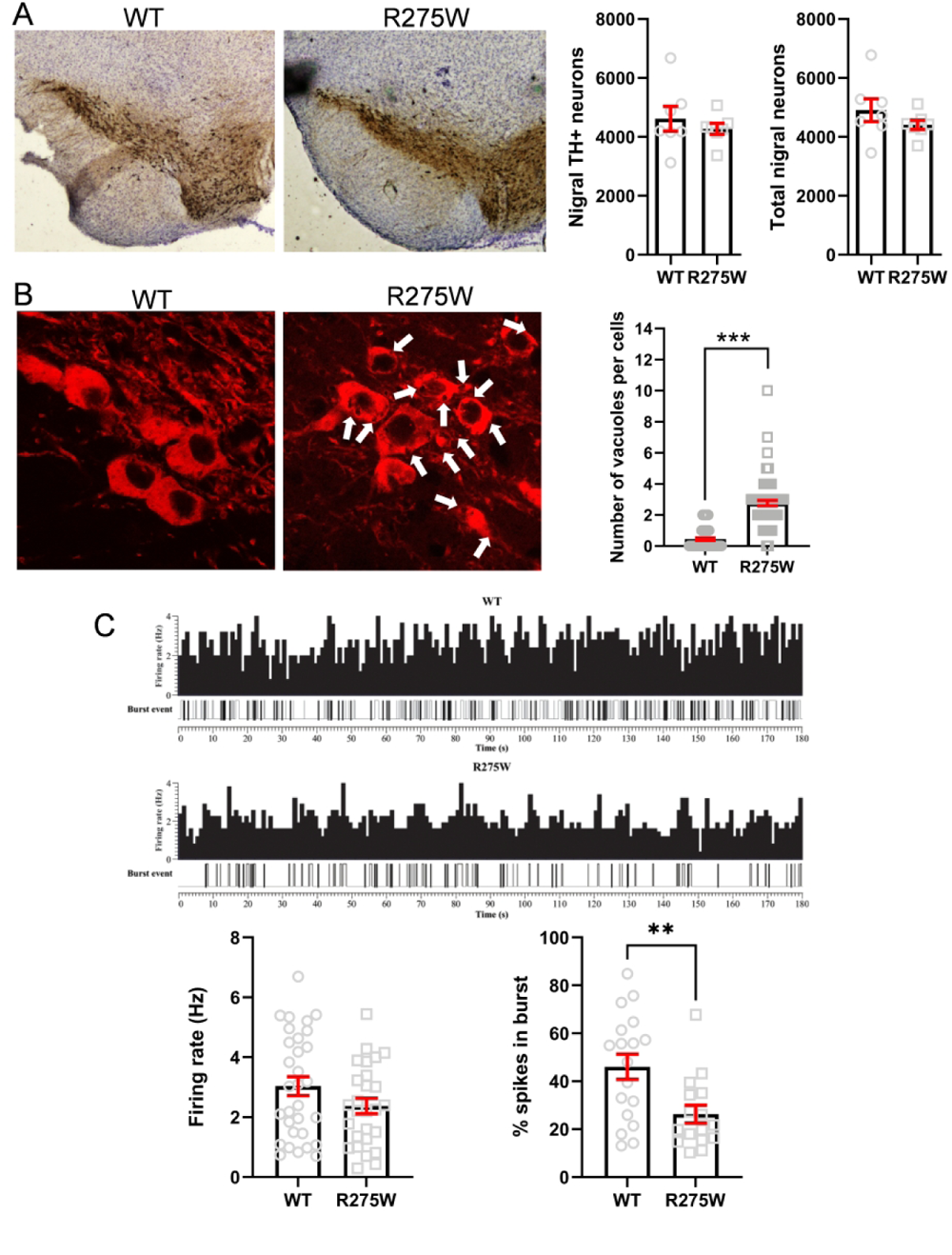
DA neuron dysfunction in young (1-month-old) *Prkn*R275W mice. **A)** Representative images showing TH immunoperoxidase labelling in the SNc of WT and *Prkn*R275W mice at 1 month of age. DA neuron quantification performed by unbiased stereology counting of TH+ neurons and Nissl+ neurons is shown on the right. At 1 month of age, the number of SNc DA neurons was similar in WT and *Prkn*R275W mice (WT 4618 ± 419 *vs*. R275W 4276 ± 190, n = 7 mice analysed for each genotype. Nissl+ neurons WT 4903 ± 386 *vs*. R275W 4407 ± 159, n = 7 mice analysed for each genotype; two-tailed unpaired t-test; p > 0.05). **B)** Representative images showing DA neurons (TH labelling in red) in the SNc of WT and *Prkn*R275W mice at 1 month of age. SNc neurons of *Prkn*R275W mice showed cytoplasmic vacuolization. The white arrows indicate the vacuoles in the cytoplasm of DA neurons. The graphs on the right show the number of vacuoles per cell (n = 75 cells analyzed for each genotype, 3 mice for each genotype, Mann– Whitney test, ***p < 0.001). **(C)** Representative histograms of spontaneous single-neuron firing rate activity in WT and *Prkn*R275W mice at 1 month of age. The data quantification is represented in the histograms below the figure. The tonic firing activity tended to be lower in *Prkn*R275W mice than in WT mice, but the difference did not reach statistical significance (WT 3.04 ± 0.32 Hz *vs*. *Prkn*R275W 2.37 ± 0.26 Hz, n = 31 SNc DA neurons from 8 WT mice, n = 28 neurons from 7 *Prkn*R275W mice, p = 0.117 unpaired Student’s t-test). The percentage of spikes in burst was significantly lower in R275W than in WT mice (WT 46.0 ± 5.2% *vs*. *Prkn*R275W 26.3 ± 3.7%; **p = 0.0053, t = 2.996, df = 32, unpaired Student’s t-test).

To confirm dysfunctional SNc DA neurons, *in-vivo* recording of the spontaneous activity of SNc DA neurons was carried out in 1-month-old *Prkn*R275W mice. The tonic firing activity tended to be lower in *Prkn*R275W mice than in WT mice, but the difference did not reach statistical significance (**Fig. 2C** **and** **Fig. S4**). However, the percentage of spikes that occurred in bursts was lower in *Prkn*R275W mice than in WT controls (**Fig. 2C**). Overall, these results point to early pathophysiological changes in SNc DA neurons of *Prkn*R275W mice.

### DA neuron loss, decreased striatal DA content, decreased striatal DA release, and motor impairment in adult (6-12-month-old) *Prkn*R275W mice

To test the hypothesis that the changes in DA neuron morphology and function observed in young mice precede DA neuron death, we counted the number of TH+ neurons in the SNc of adult WT and *Prkn*R275W mice. At 6 and 12 months of age, *Prkn*R275W mice showed a 23% and 28% decrease in DA neurons, respectively (**Fig. 3A**). The decrease in DA neurons was confirmed by stereological count of Nissl+ neurons (**Fig. 3A**). Cytoplasmic vacuolizations was confirmed in DA neurons of 6-month-old *Prkn*R275W mice (**Fig. S5**). Since recent studies showed that SOX6 expression distinguishes DA neuron populations more vulnerable to PD-related degeneration in the SNc,^17,18^ we also analyzed the number of SNc SOX6+ neurons. At 12 months of age, *Prkn*R275W mice showed a statistically significant reduction of double SOX6+/TH+ neurons in the SNc, from 39% in WT mice to 21% in *Prkn*R275W mice (**Fig. S6**).

**Figure 3.**
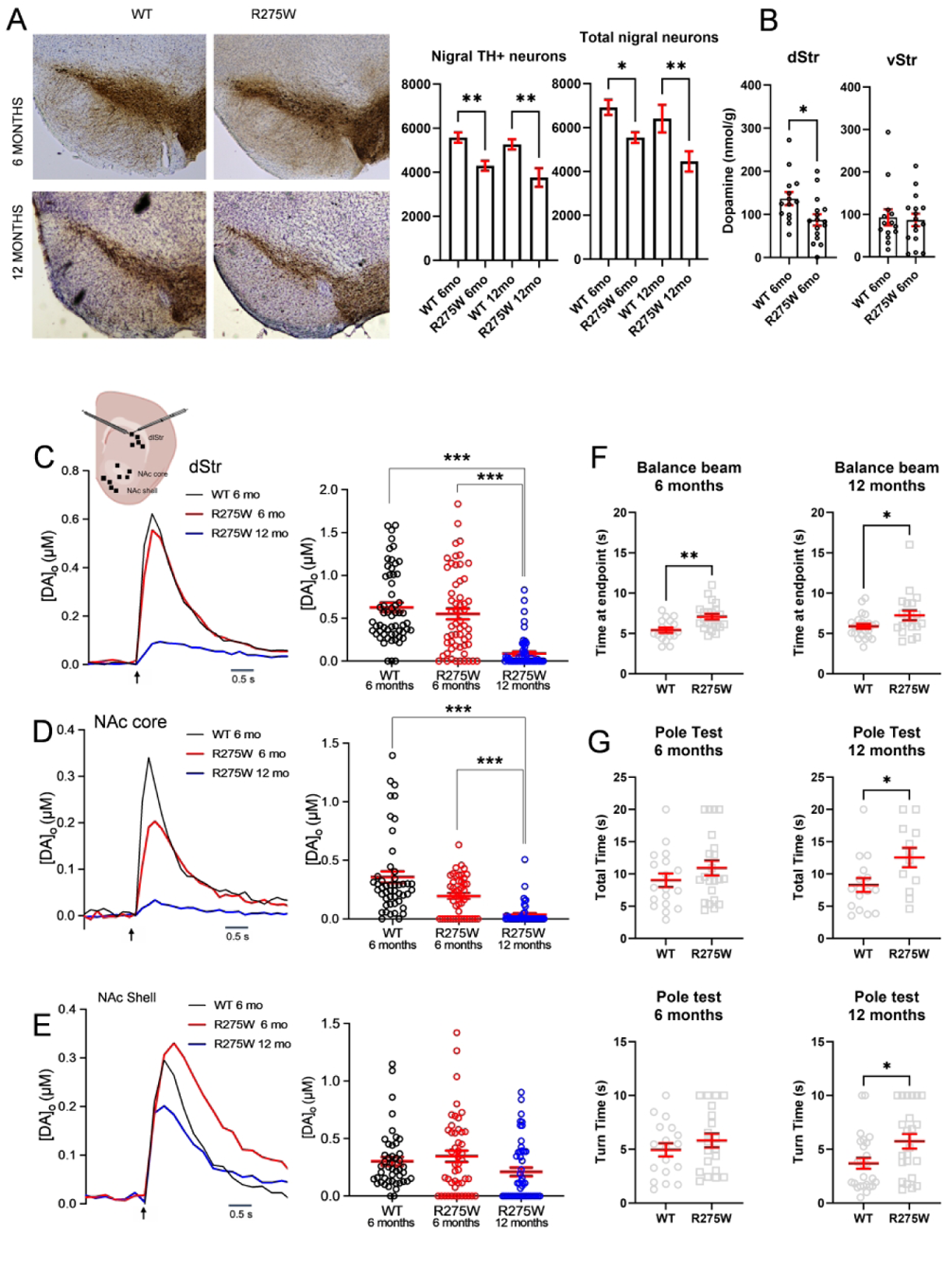
DA neuron loss, decreased striatal DA content and release, and motor impairment in adult (6-12-month-old) ParkinR275W mice. **(A)** Representative images showing TH immunoperoxidase labelling in the SNc of WT and *Prkn*R275W mice at 6 and 12 months of age. *Prkn*R275W mice showed a reduction in TH staining. The graphs on the right show TH+ neuron quantification performed by unbiased stereology (6 mo WT 5578 ± 235 *vs.* 6 mo R275W 4298 ± 228, **p = 0.0055, n = 8 WT and n = 10 R275W mice; 12 mo WT 5266 ± 231 *vs.* 12 mo R275W 3764 ± 417, **p = 0.0029, n = 7 WT and n = 8 R275W mice, one-way ANOVA followed by Šídák’s multiple comparisons test) and the number of Nissl+ neurons (6 mo WT 6919 ± 350 *vs.* 6mo R275W 5543 ± 242, *p = 0.0426, n = 8 WT and n = 10 R275W; 12 mo WT 6404 ± 635 *vs*. 12 mo R275W 4457 ± 461, **p = 0.0079, n = 7 WT and n = 8 R275W mice, one-way ANOVA followed by Šídák’s test). **(B)** Representative histograms of DA tissue content in dorsal striatum (dStr) and ventral striatum (vStr) of WT and *Prkn*R275W mice at 6 months of age. *Prkn*R275W mice showed a reduction of tissue DA content in dStr (nmol/g: 6 mo WT 136.7 ± 15.2 *vs.* 6 mo R275W 87.2 ± 13.2, t = 2.464, df = 28, n = 14-16 samples from 6 mice per genotype, unpaired t-test, *p = 0.0201), whereas no difference was seen in vStr (6 mo WT 93.1 ± 19.1 nmol/g *vs.* 6 mo *Prkn*R275W 87.0 ± 14.8 nmol/g, p = 0.8861, unpaired U-test; n = 14-16 samples from 6 mice per genotype, p > 0.05). **(C)** Upper part: Representation of section of mouse brain showing typical level of forebrain slices used to study axonal DA release (modified from Franklin and Paxinos, 2008). At this level, local electrical stimulation can be used to evoke DA release in dlStr and in NAc core and shell in the same *ex vivo* slice. The curve shows the average evoked [DA]_o_ (single-pulse stimulation) in dlStr. Arrow indicates time of stimulation. In 6-month-old mice, evoked [DA]_o_ did not differ between the two genotypes (6 mo WT 0.63 ± 0.05 µM *vs.* 6 mo R275W: 0.55 ± 0.06 µM, p = 0.1026, Kruskal-Wallis with Dunn’s test; n = 59-57 release records from 6 mice per genotype). Evoked [DA]_o_ differed between dlStr of 12-month-old R275W mice and 6-month-old mice of either genotype (12 mo *Prkn*R275W: 0.09 ± 0.02 µM, ***p < 0.001 *vs*. 6 mo WT; ***p < 0.001 *vs*. 6 mo R275W; Kruskal-Wallis with Dunn’s test; n = 57-58 release records from 6 mice per genotype). **(D)** In NAc core, average evoked [DA]_o_ (single pulse stimulation) did not significantly differ between the two genotypes at 6 months of age (6 mo WT: 0.36 ± 0.05 µM *vs.* 6 mo R275W: 0.20 ± 0.03 µM, Kruskal-Wallis with Dunn’s test, p = 0.063). In NAc core of 12-month-old R275W mice evoked [DA]_o_ was lower as compared to 6-month-old mice of both genotypes (12 mo R275W 0.035 ± 0.013 µM, ***p < 0.001 *vs*. 6 mo WT; ***p < 0.001 *vs*. 6 mo R275W; Kruskal-Wallis with Dunn’s test; n = 46-48 release records from 6 mice per genotype)**. E)** In NAc shell, average evoked [DA]_o_ (100 Hz 5-pulse stimulation) did not differ between the two genotypes at 6 months of age (6 mo WT: 0.30 ± 0.04 µM *vs.* 6 mo *Prkn*R275W: 0.35 ± 0.05 µM, p > 0.05). In 12 month-old *Prkn*R275W mice average evoked [DA]_o_ was similar to that measured in 6-month-old mice of both genotypes (12 mo *Prkn*R275W 0.21 ± 0.04 µM, p > 0.05 *vs*. 6 mo WT; p > 0.05 *vs*. 6 mo R275W). **(F)** Six and 12 month-old parkinR275W mice were significantly impaired in the balance beam test (6 mo WT 5.4 ± 0.3 *vs.* 6 mo *Prkn*R275W 7.1 ± 0.3, unpaired t-test, t = 3.456, df = 37, **p = 0.0014, data from 17 WT and 22 *Prkn*R275W mice; 12 mo WT 6.0 ± 1.1 s *vs.* 12 mo R275W 12.6 ± 1.5 s, unpaired t-test, t = 1.836, df = 41, *p = 0.0431, data from 23 WT and 20 R275W mice). **(G)** Results from pole test were similar between WT and *Prkn*R275W mice at 6 months of age, whereas 12-month-old R275W mice showed an impairment as compared to WT mice of the same age (total time pole test 12 mo WT 8.3 ± 0.3 s *vs.* 12 mo *Prkn*R275W 7.2 ± 0.6, unpaired t-test, t = 2.367, df = 26, *p = 0.0256; turn time pole test 12 mo WT 3.7 ± 0.5 s *vs.* 12 mo R275W 5.7 ± 0.7 s, unpaired t-test, t = 2.462, df = 47, *p =0.0176, data from 23 WT and 20 R275W mice).

These results demonstrate that the *Prkn*R275W transgenic strain recapitulates the early degeneration of nigrostriatal DA neurons typical of ARJP patients.^2–4^

To assess whether DA neuron degeneration resulted in decreased tissue DA levels in target regions, we determined DA content in dorsal striatum (dStr) and ventral striatum (vStr) from 6-month-old *Prkn*R275W and WT mice using high-performance liquid chromatography (HPLC). Consistent with the significant decrease in SNc DA neuron number in *Prkn*R275W *vs*. WT mice at 6 months, total tissue DA content in the dStr was significantly lower in *Prkn*R275W mice compared to that in WT mice (**Fig. 3B**). No difference was observed in DA content of the vStr, which contains nucleus accumbens (NAc) core and shell (**Fig. 3B**).

We also examined the influence of the *Prkn*R275W mutation on dynamic DA release in *ex vivo* corticostriatal slices from 6-month-old and 12-month-old mice. Fast-scan cyclic voltammetry (FSCV) was used to quantify evoked [DA]_o_ in the dorsolateral striatum (dlStr) and NAc core and shell, with sampling of 3-5 sites in each region for every slice tested (**Fig. 3C, inset**). Evaluation of evoked [DA]_o_ in the dlStr of slices from *Prkn*R275W and WT mice recorded under control conditions indicated no difference in [DA]_o_ amplitude between genotypes at 6 months of age. At 12 months, evoked [DA]_o_ in dlStr of R275W mice was significantly lower than that in 6-month-old WT and *Prkn*R275W mice (**Fig. 3C**). Evoked [DA]_o_ in the NAc core from 12-month-old *Prkn*R275W mice was lower than in the core from WT or *Prkn*R275W mice at 6 months (**Fig. 3D**). In the Nac shell, we found no difference in evoked [DA]_o_ between *Prkn*R275W and WT mice (**Fig. 3E**).

To analyze the consequences of DA neuron loss and striatal DA depletion in 6-12-month-old *Prkn*R275W mice, we tested whether these mice also displayed motor dysfunction. Six and 12-month-old *Prkn*R275W mice were significantly impaired in the balance beam test (**Fig. 3F**). In the pole test, performances were similar between WT and *Prkn*R275W mice at 6 months of age whereas at 12 months of age *Prkn*R275W were impaired (**Fig. 3G**). Finally, in the rotarod test, 6 and 12-month-old *Prkn*R275W mice performed similarly to their WT counterparts (**Fig. S7**). These data indicate that *Prkn*R275W mice have deficits in fine balance and motor coordination already by 6 months of age.

### DA neuron loss and overt motor impairment in aged (18-month-old) *Prkn*R275W mice

At 18 months of age, *Prkn*R275W mice showed a 40% decrease in the number of DA neurons (**Fig. 4A**). To analyze the consequences of progressive DA neuron loss on motor function, we examined motor behavior. Eighteen-month-old *Prkn*R275W mice performed significantly worse than WT mice in the rotarod test and balance beam test (**Fig. 4B**). Pole test results are not shown because 18 month-old mice were too impaired to accomplish the task. Overall, these data confirm that *Prkn*R275W mice display progressive, age-dependent motor impairments.

**Figure 4.**
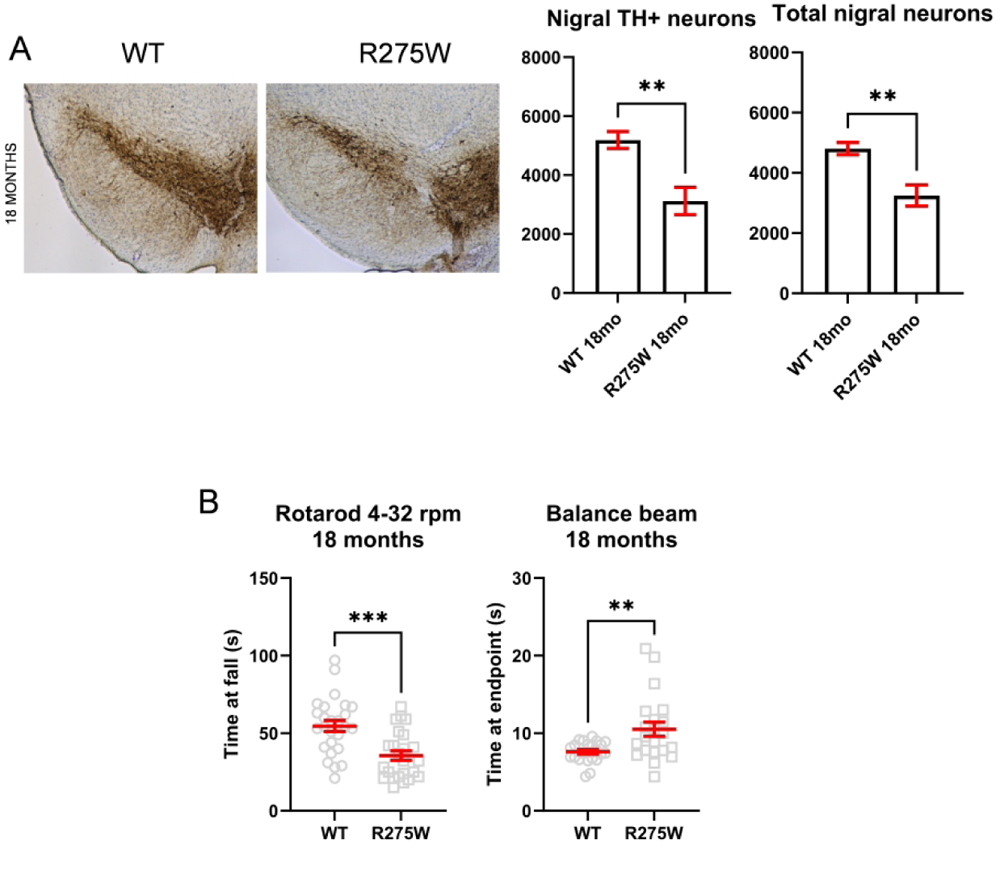
DA neuron loss and impairment in the motor balance and coordination in aged (18-month-old) *Prkn*R275W mice. **(A)** Representative images showing TH immunoperoxidase labelling in the SNc of WT and *Prkn*R275W mice at 18 months of age. *Prkn*R275W mice showed a reduction in the levels of staining. The graphs on the right show the DA neuron quantification performed by unbiased stereology by analyzing the number of TH+ neurons (18 mo WT 5190 ± 285 *vs.* 18 mo R275W 3120 ± 464, unpaired t-test, t=3.804, df=10, **p = 0.0035, data from 6 WT and 6 R275W mice). The progressive decrease in DA neurons was confirmed by stereological count of Nissl positive neurons in SNc (18 mo WT 4805 ± 204 *vs.* 16 mo R275W 3245 ± 349, unpaired t-test, t=3.861, df=16, p = **0.0014, data from 9 WT and 9 *Prkn*R275W mice). **(B)** The graph shows the performance at rotarod and balance beam test of 18-month-old mice. Eighteen-month-old R275W mice performed significantly worse in both the rotarod and the balance beam test (rotarod test: 18 mo WT 54.6 ± 3.6 s *vs.* 18 mo *Prkn*R275W 35.5 ± 3.2 s, unpaired t-test, ***p = 0.0002, data from 26 WT and 24 R275W mice; balance beam: 18mo WT 7.6 ± 0.3 s *vs.* 18 mo R275W 10.6 ± 0.9 s, unpaired t-test with Welch’s correction, **p = 0.0063, data from 22 WT and 21 R275W mice). Pole test results are not shown because 18-month-old *PrknR275W* mice were too impaired to perform the task.

## Discussion

Many pathogenic variants in *PRKN* gene, including exonic rearrangements and missense, nonsense, and frameshift mutations, cause juvenile parkinsonism. The present work describes the phenotype of the knock-in mouse model expressing the PARKIN pathogenic variant p.Arg275Trp (R275W). This mutation was chosen because it is the missense mutation with the highest allelic frequency in *PRKN* patients and has been identified in many ARJP patients as either compound heterozygous or homozygous mutation (https://www.mdsgene.org).

There are no specific clinical features that distinguish among patients carrying the various pathogenic variants in the *PRKN* gene. The lack of genotype-phenotype association reflects the fact that all the pathogenic variants in the *PRKN* gene cause protein loss-of-function or absence of protein.^14,19^ Consistent with these findings, we found a 70% decrease in PARKIN protein levels in the brain of *Prkn*R275W mice. This also agrees with previous studies showing protein conformational alteration and loss-of-function phenotype for R275W patients and for most *PRKN* mutations.^14,15,19^ This downregulation of endogenous PARKIN levels associated with the presence of the missense *Prkn*R275W mutation could be the reason why this model exhibits a parkinsonian phenotype, whereas *Prkn* KO mouse models do not. Indeed, genetic compensation is a feature of KO models but not of knockdown and mutant models.^11^ We can hypothesize that low expression levels of the PARKIN pathogenic variant R275W cause the phenotype without inducing genetic compensation. This hypothesis could be confirmed by future studies in mouse models carrying other pathogenic missense mutations in *Prkn* gene.

The neuropathology of patients carrying *PRKN* mutations is characterized by a loss of nigrostriatal DA neurons not associated with alpha-synuclein neuronal inclusions.^2^ By performing stereological counts of TH+ and total neurons in SNc, we report a loss of DA neurons in the *Prkn*R275W mouse starting at 6 months of age. Considering that 6 months of murine age correspond to ∼30 years of age in humans,^20^ the 23% loss of DA neurons in 6-month-old *Prkn*R275W mice correlates well with the median age at symptom onset in humans (31 years).^3,4^

In 12-month-old and 18-month-old *Prkn*R275W mice, we found a loss of 28% and 40% respectively of TH+ neurons, which correlates with the slow progression of symptoms seen in ARJP patients.^3,4,21^ A further confirmation of the DA neuron loss comes from the labeling of substantia nigra for SOX6, a recently discovered marker of DA neuron vulnerability in PD. ^17,18^

The loss of DA neurons is also in agreement with the decrease of total DA content in the dlStr seen as early as at 6 months of age in *Prkn*R275W mice. Interestingly, at 6 months, DA release assessed by FSCV was relatively preserved in *Prkn*R275W mice, even in the dlStr which showed a significant decrease in DA content. This finding is consistent with previous studies in 6-OHDA lesioned rats, showing relatively constant peak evoked [DA]_o_ *in vivo* in spite of progressively larger DA loss.^22^ This process, which Bergstrom and Garris called passive stabilization, reflects loss of DA uptake sites as well as of release sites, leading to maintenance of net evoked [DA]_o_. This maintenance of dynamic DA release levels is also a likely contributing factor to the relative preservation of motor function seen in 6-month-old *Prkn*R275W mice. Other compensatory mechanisms might however also play a role. For example, DA release evoked by local electrical stimulation is triggered not only by the depolarizing stimulus, but also by concurrently released acetylcholine (ACh) from striatal cholinergic interneurons. In fact, endogenous ACh acts at nicotinic ACh receptors (nAChRs) on DA axons to promote release.^23–26^ Therefore, an additional contributing factor to preserved evoked [DA]_o_ in 6-month-old *Prkn*R275W mice might be local boosting of DA release by ACh acting at nAChRs, a hypothesis which will be tested in future studies.

Strikingly, in the striatum of *Prkn*R275W mice evoked [DA]_o_ was significantly lower at 12 months of age than at 6 months. Consistent with the progressive loss of DA neurons and striatal DA signaling, motor behavior showed progressive impairment between 6 and 18 months, as indicated by increasingly poor performance in rotarod, balance beam and pole tests.

Finally, it is interesting to note that in the *Prkn*R275W model histological and functional alterations, including cytoplasmic vacuolization and DA neuron firing changes, already appear at 1 month of age, i.e. long before the loss of DA neurons. Since cytoplasmic vacuolization is a morphological phenomenon that often accompanies cell dysfunction and precedes cell death,^27^ these findings reveal early pathology, consistent with observations in both human and mouse DA neurons expressing PARKIN mutations.^16^ Since PARKIN is concentrated at presynapses^28^ and associates with the cytoplasmic surface of the synaptic vesicle,^29^ *PRKN* mutations may directly dysregulate DA release. Thus, early DA neuron firing changes are compatible with the proposed synaptic function of parkin.^30^

In conclusion, the *Prkn*R275W mouse recapitulates many of the key features of ARJP, including DA neuron loss in the midbrain, DA depletion in the striatum, and progressive motor impairment. We therefore propose that the *Prkn*R275W mouse is an extremely useful tool for the study of the causative mechanisms of PD and the development of neuroprotective therapies for this disabling disease.

## Supplementary material

Supplementary material is available at Brain online.

### Abbreviations

β2M: beta-2 microglobulin
ACh: acetylcholine
ARJP: autosomal recessive juvenile parkinsonism
DA: dopamine
dlStr: dorsolateral striatum
FSCV: fast-scan cyclic voltammetry
HPLC: High-Performance Liquid Chromatography
KO: knockout
nAChRs: acetylcholine receptors
NAc: nucleus accumbens
PD: Parkinson’s disease
SNc: substantia nigra pars compacta
TH: tyrosine hydroxylase
vStr: ventral striatum
VTA: ventral tegmental area
WT: wild-type

## Supplementary Materials

### ABBREVIATIONS SUPPLEMENTARY TEXT

β2M: beta-2 microglobulin
aCSF: artificial cerebrospinal fluid
BSA: bovine serum albumin
DAPI: 4′,6-diamidino-2-phenylindole
DDC: dopa decarboxylase
dlStr: dorsolateral striatum
FSCV: fast-scan cyclic voltammetry
HPLC: high-performance liquid chromatography
NAc: nucleus accumbens
OD: optical density
PBS: phosphate buffered saline
PBST: PBS containing 0.3% Triton X-100
PFA: paraformaldehyde
PSB: pontamine sky blue
qPCR: quantitative real-time PCR
RT: reverse transcriptase
SDHA: succinate dehydrogenase complex flavoprotein subunit A
SDS: sodium dodecyl sulfate
sgRNA: single guide RNA
TBS-T: tris-buffered saline containing 0.05% tween-20
TH+: TH positive
VDAC: voltage-dependent anion channels

## Material and Methods

### Constitutive knock-in of a point mutation in the *Prkn* gene via CRISPR/Cas9-mediated gene editing

Constitutive knock-in of a point mutation in the *Prkn* gene via CRISPR/Cas9-mediated gene editing was performed by Taconic Biosciences. The targeting strategy was based on NCBI transcript NM_016694_3, which corresponds to Ensembl transcript ENSMUST000000191124. The human R275 in the *Prkn* mouse orthologue (Ensembl gene ID: ENSMUSG00000023826), corresponds to the amino acid R274. The R274W mutation was introduced into exon 7 using specific single guide RNA (sgRNA) and oligonucleotide. The Cas9 mRNA, the specific sgRNA and oligonucleotide were injected into C57BL/6NTac zygotes. Alongside the exchange of the first nucleotide on the R274 codon, two additional silent mutations were introduced close to the site of the mutation to create a novel AflIII restriction site that allows for genotyping of the mutated allele. C57BL/6NTac-R275W mice were backcrossed to C57BL/6 mice (Charles River) for at least 10 generations before starting the phenotypic characterization.

### Animals

All experiments were conducted with the goal of minimizing the number of sacrificed animals according to the principle of the 3Rs (replacement, reduction and refinement). Sex-matched C57BL/6 WT mice were used as controls. Since *PRKN*-related ARJP is not sex-linked, both male and female mice (in equal ratio) were used, minimizing the numbers of animals of a given sex required. Mice were maintained and bred at the animal house of Ospedale San Raffaele in compliance with institutional guidelines and international laws (EU Directive 2010/63/EU EEC Council Directive 86/609, OJL 358, 1, December 12, 1987, NIH Guide for the Care and Use of Laboratory Animals, U.S. National Research Council, 1996), and also housed at the NYU Grossman School of Medicine in accordance with NIH guidelines and with the approval of the local Institutional Animal Care and Use Committee. Animals were housed in a climate-controlled room with a 12-hour light/dark cycle. Mice were given unlimited access to food and water.

### RNA extraction and quantitative real-time polymerase chain reaction

Total brain RNA from 1-month-old WT and *Prkn*R275W mice was purified using TRIzol™ Reagent (Invitrogen) according to the manufacturer’s instructions. Briefly, brains were homogenized in 1.5 mL of TRIzol then centrifuged for 5 min at 4°C and 12,000 x *g*. The clear supernatant was transferred to a new tube and incubated for 5 min at room temperature. Then, 300 µL of chloroform was added and the samples were mixed by shaking and centrifuged at 4°C and 12,000 x *g* for 15 min. The aqueous phase containing RNA was removed and incubated with 500 µL of isopropanol for 10 min at 4°C. After centrifugation at 4°C and 12,000 x *g* for 10 min, the RNA pellet was washed with 1 mL of 75% ethanol and centrifuged again at 4°C and 7,500 x *g* for 5 min. Finally, RNA was resuspended in RNAase-free water and quantified using NanoDrop (ThermoFisher).

Reverse transcription was performed with ProtoScript^®^ II Reverse Transcriptase (RT) (New England Biolabs) according to the manufacturer’s instructions. Briefly, 1 μg of RNA was incubated with 10 mM dNTPs (Promega) and 63 µM Random Hexamers (Promega) at 65°C for 5 min. Each sample was then incubated with ProtoScript II Buffer, 0.1 M dithiothreitol (DTT), and ProtoScript II RT (200 U/μL) at 25°C for 5 min, 42°C for 60 min and 65°C for 20 min. The cDNA obtained served as a template for quantitative real-time PCR (qPCR) based on the TaqMan methodology (Life Technologies).

For Taqman probe-based qPCR, the TaqMan™ Gene Expression Master Mix (Applied Biosystems™) and the Taqman Assay (ThermoFisher) were combined with 100 ng of cDNA. All reactions were performed and quantified using the CFX96 touch real-time PCR detection system (Bio-Rad Laboratories). Cycling parameters were 50°C for 2 min, 95°C for 10 min, then 40 cycles at 95°C for 15 s and 60°C for 60 s.

Results were calculated by the 2^−ΔΔCT^ method and are presented as percent change in gene expression normalized to the endogenous reference gene *Beta-2 microglobulin* (β2M) and relative to the WT group. TaqMan Assay used are: *Prkn* Mm00450187_m1, β2M Mm00437762_m1.

### Western blotting

Brain tissues were homogenized in ice-cold lysis buffer (50 mM Tris-HCl pH 7.5, 150 mM NaCl, 1 mM EDTA, 1% NP-40, 0.5% sodium deoxycholate, 0.1% Sodium dodecyl sulfate-SDS) added with Complete Protease Inhibitor Cocktail (Sigma). Lysates were incubated in ice for 30 min, sonicated in ice for 15 s with the ultrasonic processor UP100H (Hielscher) at a frequency of 30 kHz and a power output of 80%, then centrifuged at 12,000 x *g* at 4°C for 20 min. Proteins in the supernatant were quantified by BCA assay (Thermo Scientific) according to the manufacturer’s instructions. Samples were diluted with 3X SDS loading buffer (188 mM Tris-HCl, pH 6.8, 6% SDS, 30% glycerol, 0.3% bromophenol blue) and 1 M 1,4-Dithiothreitol (DTT) (final concentration of DTT was 100 mM) and heated to 95°C for 10 min. Protein extracts (20 μg) were separated on NuPAGE 4-12% Bis-Tris gels (Life Technologies) with NuPAGE MOPS SDS Running Buffer (Life Technologies) and the MagicMark XP Standard molecular weight marker (Life technologies). After electrophoresis, gels were transferred onto nitrocellulose membranes (GE Healthcare) for 2 h at 400 mA constant current at 4°C in transfer buffer (25 mM Tris, 192 mM glycine, 20% methanol). Post-transfer membranes were briefly incubated in Ponceau Solution (Sigma) to visualize proteins, rinsed in Tris-Buffered Saline containing 0.05% Tween-20 (TBS-T) and blocked with 5% non-fat dried milk in TBS-T for 1 h at room temperature and then incubated overnight at 4°C with the following primary antibodies in TBS-T: mouse anti-parkin 1:2000 (P6248, Sigma Aldrich), rabbit anti-GAPDH 1:1000 (Santa Cruz, sc-25778), rabbit anti-DDC 1:1000 (cell Signaling #13561), mouse anti-SDHA 1:10.000 (ThermoScientific #459200), mouse anti-VDAC1 1:1000 (Abcam #ab147). Membranes were then washed 3×10 min in TBS-T, incubated with horseradish peroxidase (HRP)-conjugated secondary antibodies (GE Healthcare-Amersham Biosciences) for 1 h at room temperature and then washed 3 x10 min in TBS-T. The antigens were detected using an immunodetection kit (Novex ECL, Invitrogen) according to manufacturer’s instructions. For visualization we used the Chemidoc Touch Imaging system (BioRad). Band intensities were quantified by densitometry using ImageJ software.^1^ Briefly, the images were converted to 8-bit format in order to perform uncalibrated optical density (OD). After conversion, the background was subtracted through the rolling ball radius method (default setting, ball radius value: 50.0 pixel). Each band was individually selected and circumscribed with the rectangular ROI tool selection. Data were acquired as arbitrary integrated density values. The final relative expression level was normalized to the internal standard GAPDH.

### *In Vivo* single-unit extracellular recording of SNc DA Neurons

*In vivo* single-unit extracellular recordings of SNc DA neurons were conducted according to standard protocols in the lab.^2,3^ Briefly, anesthetized mice were placed in a stereotaxic apparatus with the skull positioned horizontally. After an incision was made in the scalp, a burr hole was drilled through the skull over the SNc using stereotaxic coordinates from Paxinos and Franklin’s mouse brain atlas:^4^ 0.08–2.0 mm posterior to the interaural line and 0.7–1.7 mm lateral to midline. The spontaneous electrical activity of SNc DA neurons was recorded using single-barreled glass micropipettes at a depth from the brain surface ranging from 3.5 to 5.0 mm. The glass micropipette was filled with 2% pontamine sky blue (PSB) solution in Na-acetate (0.5 M), and had an impedance of 2-6 MΩ. Identification of recorded neurons in the SNc was based on classical electrophysiological properties of DA neurons (action potential duration >2.5 ms, a biphasic or triphasic waveform, and a slow firing rate of 0.5–10 Hz), analyzed off-line using Spike2 software (ver. 8.21; CED, UK). Burst-firing activity of these cells was also analyzed using a script for the Spike 2 software, with a burst defined as a train of at least two spikes with an initial interspike interval (ISI) ≤80 ms and a maximum ISI of 160 ms, within a regular low-frequency firing pattern and decreased amplitude from the first to the last spike within the burst.

### Stereological cell counting of SNc DA neurons

Stereological counting was performed according to published methods.^5,6^ Mice were deeply anesthetized by intraperitoneal injection of a mixture of ketamine/xylazine (100 mg/kg and 10 mg/kg, respectively; Sigma-Aldrich) and transcardially perfused with ice-cold 0.9% NaCl solution, followed by 4% paraformaldehyde (PFA) in phosphate buffered saline (PBS) (0.1 M, pH 7.4, Sigma-Aldrich). The brains were post-fixed in 4% PFA overnight at 4°C, then transferred to a 30% sucrose solution in PBS for cryoprotection and stored at −80°C. Free-floating midbrain sections (50 µm-thickness) containing the SNc (AP: −3.16 to −3.52 from bregma)^4^ were prepared using a freezing microtome. Sections were washed in PBS and incubated in 3% hydrogen peroxide/PBS for 10 min to eliminate endogenous peroxidase activity. After several washes, sections were incubated for 30 min at room temperature with blocking solution (PBS containing 0.3% Triton X-100 (PBST) and BSA 2%, and then incubated overnight at 4°C with an antibody to tyrosine hydroxylase (TH) (ab112; 1:750 in BSA 1% PBST; Abcam). Sections were then rinsed, incubated for 1 h with anti-rabbit HRP-conjugated secondary antibody (ab6721, 1:500 in BSA 1% PBST; Abcam) and TH+ neurons visualized using a DAB substrate kit (ab64238, Abcam). Sections were mounted on slides coated with gelatine, dehydrated by incubating overnight in a chloroform-alcohol mixture. For Nissl counterstaining sections were stained in cresyl violet acetate solution for 2-5 minute, rinsed in distilled water, dehydrated in graded alcohols and mounted using a xylene based mounting media (Eukitt mounting medium, Bio-Optica) and sealed with coverslips. Stereological analysis was performed by counting TH+ neurons (phenotypic marker) and cresyl violet stained cells (Nissl staining, structural marker) in the SNc. Neural cell counting was performed on 5 serial 50 µm slices that separated by 200 μm by applying an unbiased stereological sampling method based on optical fractionator stereological probe.^5,7,8^ Images were viewed at 40X magnification using a Leica DM600B motorized microscope equipped with Stereo Investigator software (MBF Europe). Neuron number is expressed as a raw value derived from the software calculation, which estimates the total number of neurons from the number of neurons within a Systematic Randomly Sampled set of unbiased virtual counting spaces covering the entire region of interest with uniform distance between unbiased virtual counting in spaces in direction X, Y and Z. Image acquisition and quantification were performed by investigators who were ‘blind’ to the experimental condition.

### Immunofluorescence

Mice were deeply anesthetized by intraperitoneal injection of a mixture of ketamine/xylazine (100 and 10 mg/kg, respectively, Sigma-Aldrich) and transcardially perfused with ice-cold 0.9% NaCl solution, followed by 4% PFA in PBS (0.1 M, pH 7.4, Sigma-Aldrich, St. Louis, MO, USA). The brains were postfixed in 4% PFA overnight at 4 °C and then transferred to a 30% sucrose solution in PBS for cryoprotection and stored at −80 °C. Coronal sections of SNc (30-μm thickness) were prepared using a freezing microtome, transferred to glass slides coated with polylysine, and dried at room temperature. Sections were then rehydrated, washed, and incubated in 10 mM sodium citrate buffer (pH 6) at 80 °C for 30 min (antigen retrieval), then kept for 20 min in ice. The slices were then treated with blocking solution, 5% normal goat serum (Sigma-Aldrich) added with 0.1% triton in PBS and incubated overnight with 1:300 anti-SOX6 (Santa Cruz, sc-393314) and 1:1000 anti-TH (Abcam, ab76442) antibodies. After three washes with PBS, the tissues were incubated for 2 h at room temperature in the dark with secondary antibodies (1:300 goat anti-mouse IgG Alexa 488 to reveal SOX6 and 1:300 goat anti-chicken IgG Alexa 465 to reveal TH, Invitrogen). After three washes in PBS, cell nuclei were counterstained with 4′,6-diamidino-2-phenylindole solution (DAPI). Slides were mounted using the Dako fluorescence mounting medium (Dako). Images were acquired using a Leica TCS SP5 confocal microscope (Leica, Wetzlar, Germany) and analyzed using ImageJ software.^9^ Both image acquisition and quantification were performed by investigators blinded to the experimental condition.

### Behavioral tests

Before proceeding to motor tests, mice were habituated to the presence of the operator by handling for 5 min per day for a minimum of 5 days prior to the first behavioral test. Since *PRKN*-related ARJP is not sex-linked, both male and female mice (in equal ratio) were used in behavioral tests. Data were analyzed to detect potential differences between males and females. No sex effect was observed in motor tests (**Fig. S8**).

#### Rotarod

Balance and coordination were assessed using a Ugo Basile rotarod apparatus (Biological Research Apparatus). The rotarod was set with increasing speed from 4 to 32 rpm with a ramp-rate of 120 seconds, and the time spent on the rotarod by each animal was recorded. A cut-off of 300 seconds was imposed. Mice were habituated to the task for four consecutive days; the fourth trial of the last test day was evaluated for statistical analysis.

#### Balance beam walking

The beam apparatus includes 1-meter beams with a flat surface of 12 mm or 6 mm located 60 cm above a table. On one end of each beam (start), a halogen lamp was placed as an aversive stimulus, and on the other end (endpoint), there was a black box filled with nesting material. The time required to traverse the beam from the start to the endpoint was recorded as the time when the anterior limbs of the animal reached the endpoint. Three trial tests were performed with 1 min rest between trials, with a final fourth trial used for statistical analysis of beam crossing speed. The beams were cleaned with 70% ethanol followed by water before placing the next mouse on the apparatus. A cut-off of 30 seconds was imposed.

#### Pole Test

In the pole test, mice were placed head facing upwards on the top of a 50 cm vertical pole with a diameter of 1.2 cm. Mice were habituated to the task in three trials over two consecutive days. On the test day, mice underwent three trials: the total time taken to turn completely and descend (latency) was recorded. A cut-off of 20 s was imposed. Data are shown as the mean of three trials per mouse performed on the test day.

#### Fast-scan cyclic voltammetry (FSCV) to quantify evoked striatal DA release in *ex vivo* slices

For brain slice preparation, each mouse was deeply anesthetized with isoflurane and the brain removed into ice-cold HEPES-buffered artificial cerebrospinal fluid (aCSF) containing in mM: 120 NaCl; 20 NaHCO_3_; 10 glucose; 6.7 HEPES acid; 5 KCl; 3.3 HEPES sodium salt; 2 CaCl_2_; and 2 MgSO_4_, equilibrated with 95% O_2_/5% CO_2_. ^10^ Coronal forebrain slices (300-μm thickness) were cut in this ice-cold solution using a Leica VT1200S vibrating blade microtome (Leica Microsystems). Slices were maintained in HEPES-buffered aCSF at room temperature for 1 h before transfer to a submersion recording chamber with 32°C aCSF flowing at 1.5 mL/min and containing in mM: 124 NaCl; 3.7 KCl; 26 NaHCO_3_; 2.4 CaCl_2_; 1.3 MgSO_4_; 1.3 KH_2_PO_4_; and 10 glucose, equilibrated with 95% O_2_/5% CO_2_. ^10–13^ Evoked DA release was monitored in the dorsolateral striatum (dlStr) and in the nucleus accumbens (NAc) core and shell using FSCV with carbon-fiber microelectrodes. Electrodes were constructed in-house from 7-μm diameter carbon fibers (Goodfellow Corporation) in pulled glass capillaries; the fibers extended 30-75 µm beyond the glass insulation and electrical contact made using Woods metal. ^14^ A Millar Voltammeter was used for FSCV; scan rate was 800 V/s and the range of the triangular voltage applied was −700 mV to +1300 mV then back to −700 mV *vs.* a Ag/AgCl reference electrode; inter-scan interval was 100 ms controlled by a Master-8 timing circuit (AMPI). Data were collected using a Digidata 1550B controlled by AxoScope 10.7 software (Molecular Devices).

Increases in extracellular DA concentration ([DA]_o_) in the dlStr, NAc core and NAc shell of WT and ParkinR275W mice at 6 months of age and Parkin2R275W mice at 12 months were evoked using a concentric microelectrode for local electrical stimulation (0.1 ms duration; 0.35 mA amplitude). Single-pulse stimulation was used in dlStr and in the NAc core, but 5-pulse stimulus trains (100 Hz) were used in NAc shell to increase the reliability of DA release detection.^12,13^ The substance detected in each region was identified as DA by the characteristic oxidation and reduction peak potentials of recorded voltammograms. ^12,14^ Two coronal slices were evaluated from each mouse; both were transferred to the recording chamber and allowed to equilibrate for 20 min before sampling was initiated. In each slice, evoked [DA]_o_ was recorded from 3-5 sites in the dlStr, NAc core and NAc shell. Multiple-site sampling was used because the site-to-site of evoked increases in [DA]_o_ in the striatum exceeds average differences between slices; recording from multiple sites across many slices minimizes sampling bias from this variability After each experiment, the electrode was calibrated in the recording chamber at 32 °C in aCSF.

### High-performance liquid chromatography (HPLC) for DA tissue content

To determine DA tissue content, the dorsal and ventral striatum were dissected from slices obtained from the same mice used for FSCV. Samples were weighed, frozen on dry ice, then stored at −80°C until analysis. Tissue content of DA was determined using high-performance liquid chromatography (HPLC) with electrochemical detection (Bioanalytical Systems), as previously reported. ^13^ For analysis, frozen samples were sonicated in ice-cold, deoxygenated eluent, centrifuged for 2 min at 13,000 x *g*, then the supernatant injected directly into the HPLC. Data are presented as nmol/g tissue wet weight.

### Statistical analysis

Data are presented as mean ± standard error of the mean (SEM). The normality test (Kolmogorov–Smirnov test) and equal variance test (Bartlett’s test) were applied. if both criteria were satisfied, two-tailed unpaired Student’s t test was used to compare two groups of data. If data were not normally distributed, Mann–Whitney test was used. If data were normally distributed but did not have equal variance, unpaired Student’s t test with Welch’s correction was used. One-way ANOVA followed by Šídák’s multiple comparisons test was used to compare more than two groups. If data were normally distributed or did not have equal variance was used to compare more than two groups. The number of DA SOX6+ neurons in WT and *Prkn*R275W mice was analysed by Fisher’s exact test. For FSCV data, normality was assessed using D’Agostino-Pearson test (α = 0.05 for normality), significance then was assessed using Kruskal-Wallis with Dunn’s test. For FSCV data, n is number of recording sites. ^11–13^ All data were analysed with Prism 8.0 (GraphPad Software Inc.).

---

**Figure S1.**
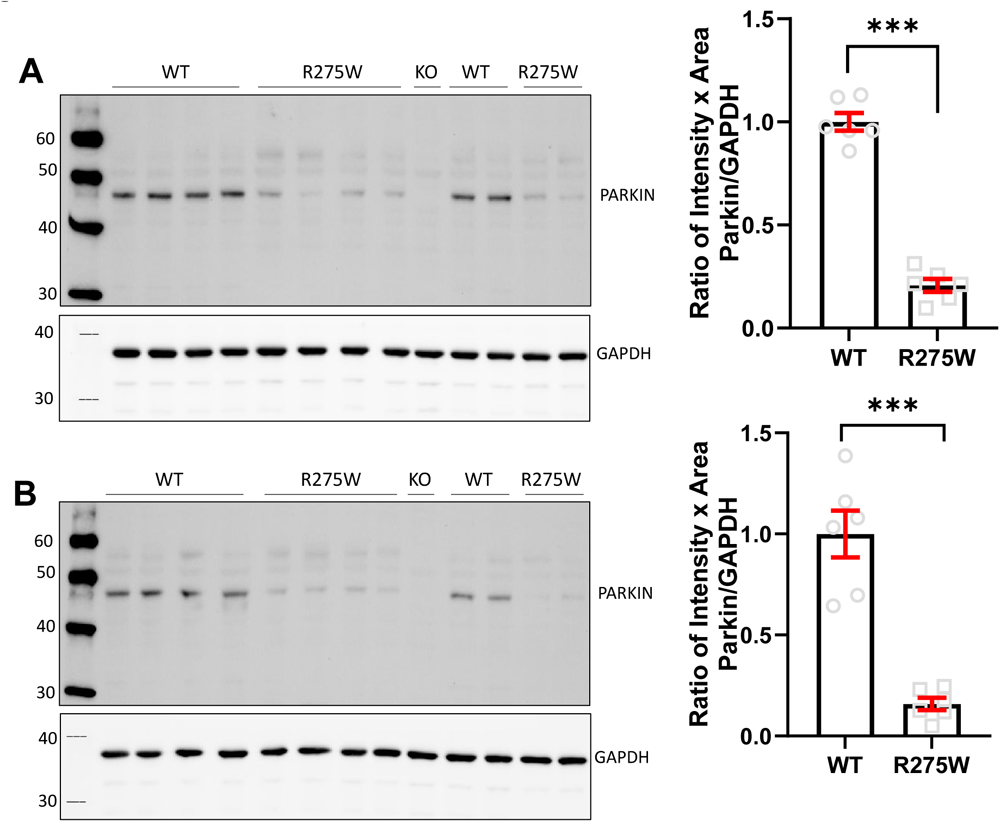
Representative western blots performed on lysates from mouse brains at 6 and 12 months of age (panel A and B, respectively) with the PARKIN monoclonal antibody Sigma P6248. *Prkn*R275W mice showed significantly reduced endogenous PARKIN protein levels when compared with WT. Histograms show mean ± SEM (6 mo WT 1.00 ± 0.04 *vs.* 6 mo *Prkn*R275W 0.21 ± 0.03, two-tailed unpaired t-test; t=14.93, df=10, ***p < 0.001, n= 6 brains from WT mice, n= 8 brains from *Prkn*R275W mice; 12 mo WT 1.00 ± 0.12 *vs.* 12 mo *Prkn*R275W 0.16 ± 0.03, two-tailed unpaired t-test; t=7.032, df=10, ***p < 0.001, n= 6 brains from WT mice, n= 8 brains from *Prkn*R275W mice). Negative control is a protein lysate from parkin KO mouse.

**Figure S2.**
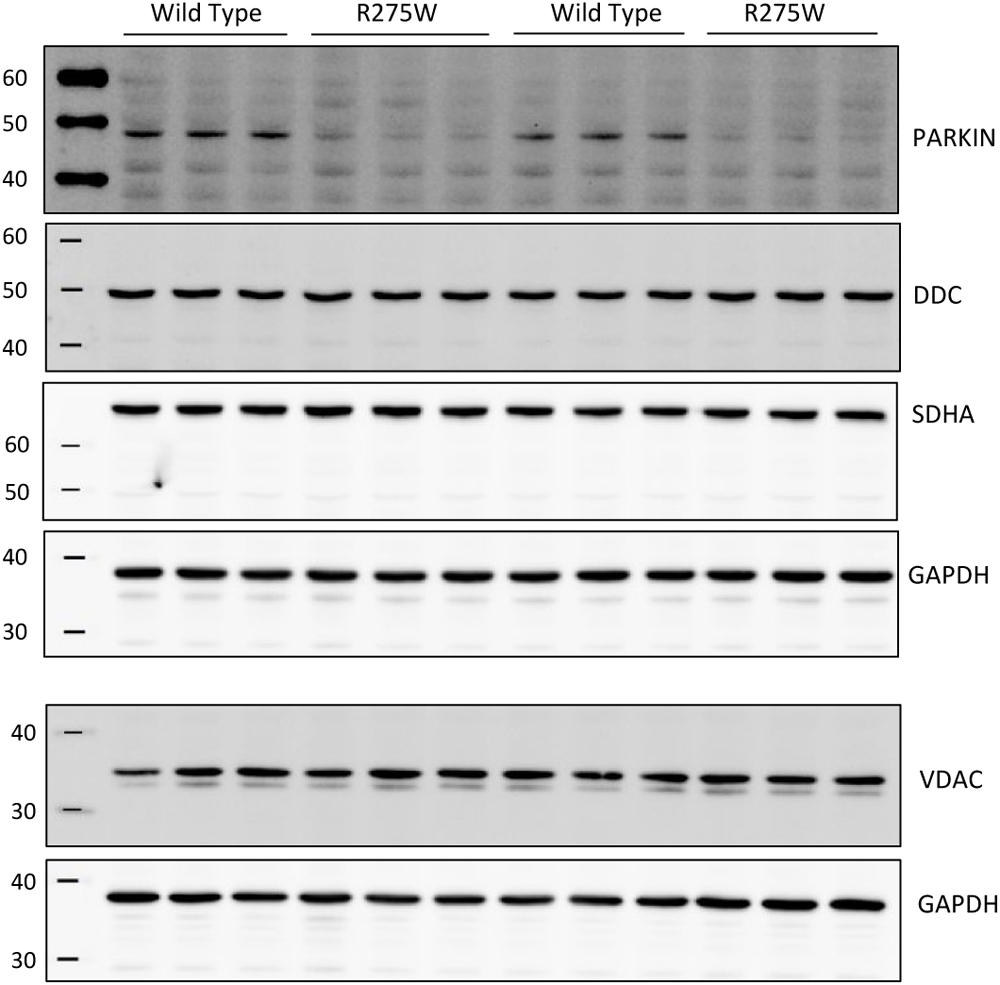
Representative western blot analysis performed with lysates prepared from total brain at 6 months of age. Parkin levels were significantly decreased in *Prkn*R275W mice, whereas the levels of other proteins, i.e. Dopa Decarboxylase (DDC), Succinate Dehydrogenase Complex Flavoprotein Subunit A (SDHA), Voltage-dependent anion channels (VDAC) are not modified. Samples were run in triplicate, with each lane loaded with lysate from an individual mouse.

**Figure S3.**
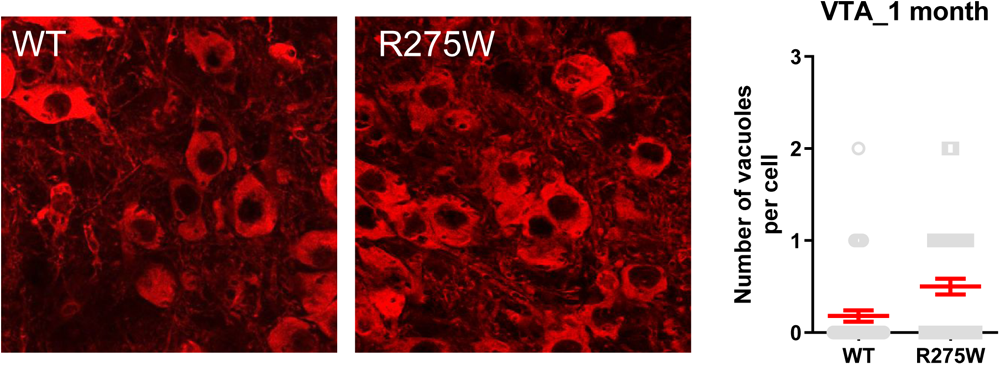
Representative images showing DA neurons (TH labelling) in the ventral tegmental area of WT and *Prk*nR275W mice at 1 month of age. The graphs on the right show the number of vacuoles per cell (n = 50 cells analyzed for each genotype, 3 mice for each genotype, Mann–Whitney test, p>0.05).

**Figure S4.**
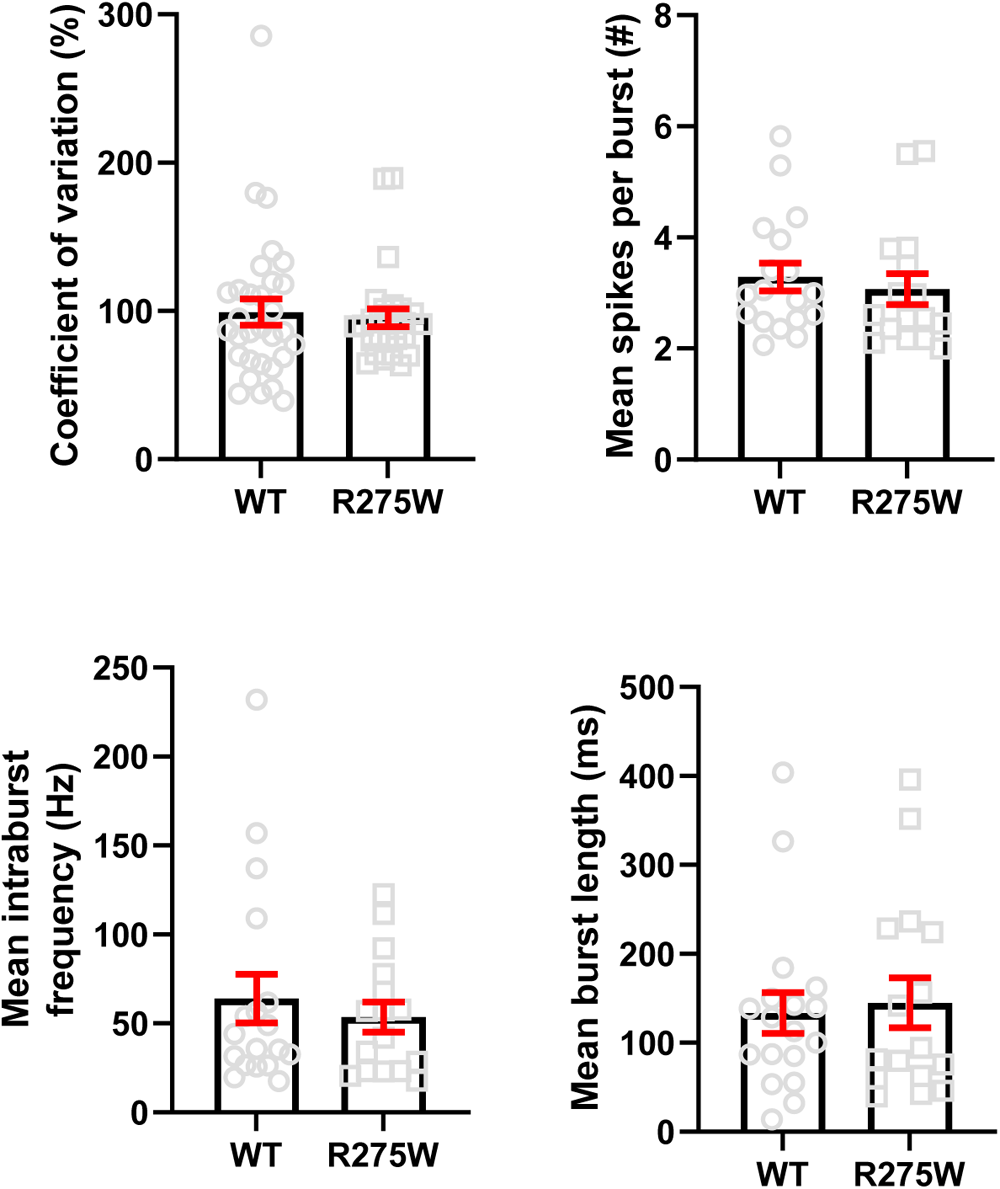
*In vivo* extracellular recording of DA neuron activity in SNc of 1-month-old WT or *Prkn*R275W mice. Coefficient of variation (COV), the number of spikes per burst, intraburst frequency and burst length were similar in WT and *Prkn*R275W mice. Data are expressed as mean ± SEM of a total of 31 neurons from 8 WT mice and 28 neurons from 7 R275W mice. p>0.05 using unpaired Student’s t-test.

**Figure S5.**
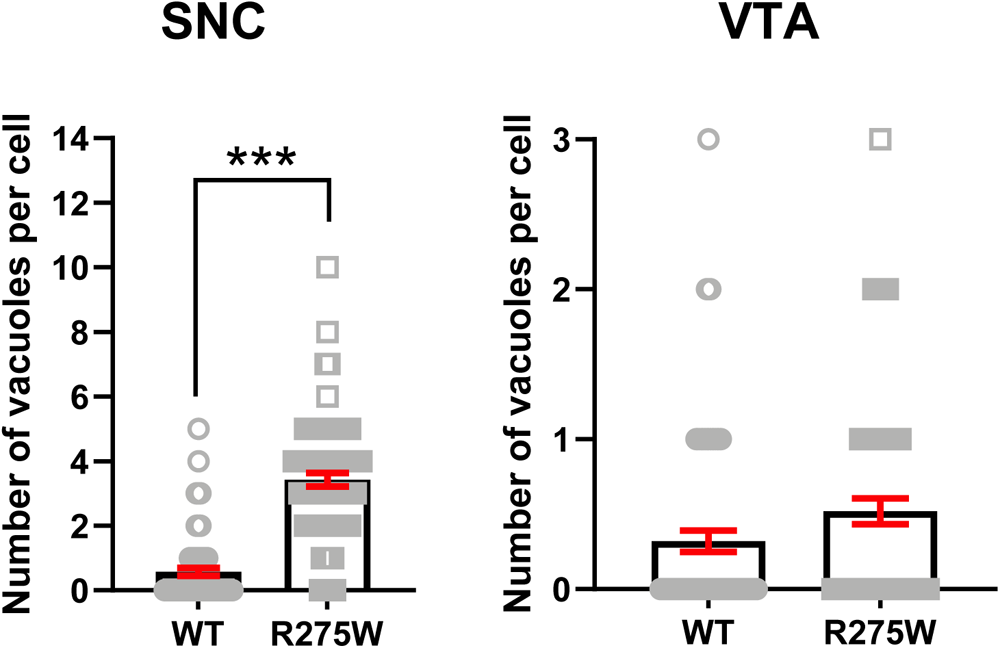
SNc neurons of PrknR275W mice at 6 months of age showed cytoplasmic vacuolization. The graphs show the number of vacuoles per cell in SNC and VTA (SNC: n = 75 cells analyzed for each genotype, 3 mice for each genotype, Mann–Whitney test, ***p < 0.001; VTA: n = 75 cells analyzed for each genotype, 3 mice for each genotype, Mann–Whitney test, p > 0.05).

**Figure S6.**
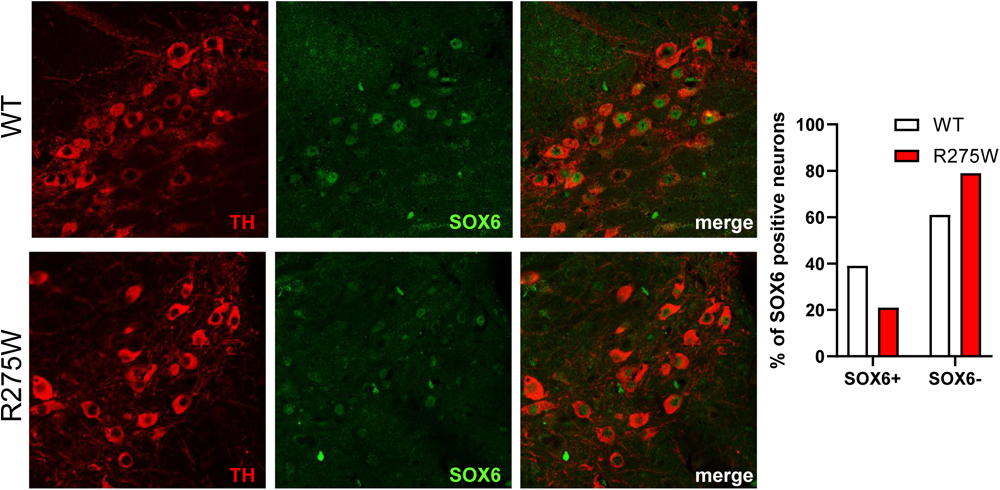
Representative images showing SOX6 immunofluorescence labelling in the SNc of WT or *Prkn*R275W mice at 12 months of age. ParkinR275W mice showed a statistically significant reduction of the percentage of SOX6+ neurons in the SNc. 39 % SNc DA neurons (TH+ cells) were SOX6+ in WT mice, whereas 21 % of SNc DA neurons were SOX6+ in *Prkn*R275W mice (Fisher’s exact test n = 401 TH+ neurons in WT section, n = 315 neurons in *Prkn*R275W sections from 3 mice for each genotype, p < 0.0001).

**Figure S7.**
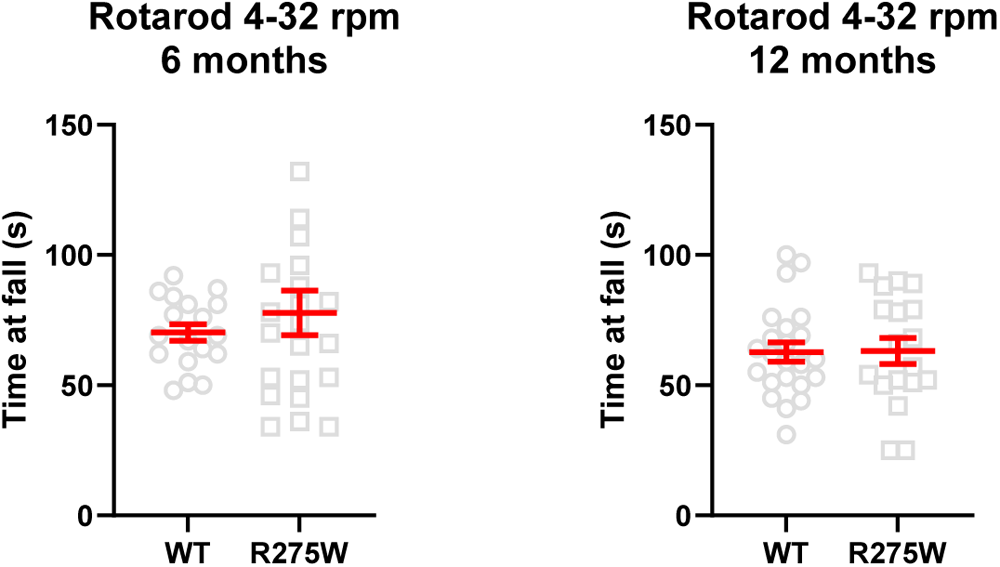
Rotarod test in 6-and 12-month-old WT and *Prkn*R275W mice. Six-and 12-month-old *Prkn*R275W mice performed similarly to WT mice (rotarod test: 6 mo WT 70.3 ± 3.2 s *vs.* 6 mo *Prkn*R275W 77.8 ± 8.5 s, unpaired t-test p > 0.05, data from 18 WT and 22 *Prkn*R275W mice; 12 mo WT 62.7 ± 3.6 s *vs.* 12 mo *Prkn*R275W 63.2 ± 5.0 s, unpaired t-test p >0.05, data from 31 WT and 25 R275W mice).

**Figure S8.**
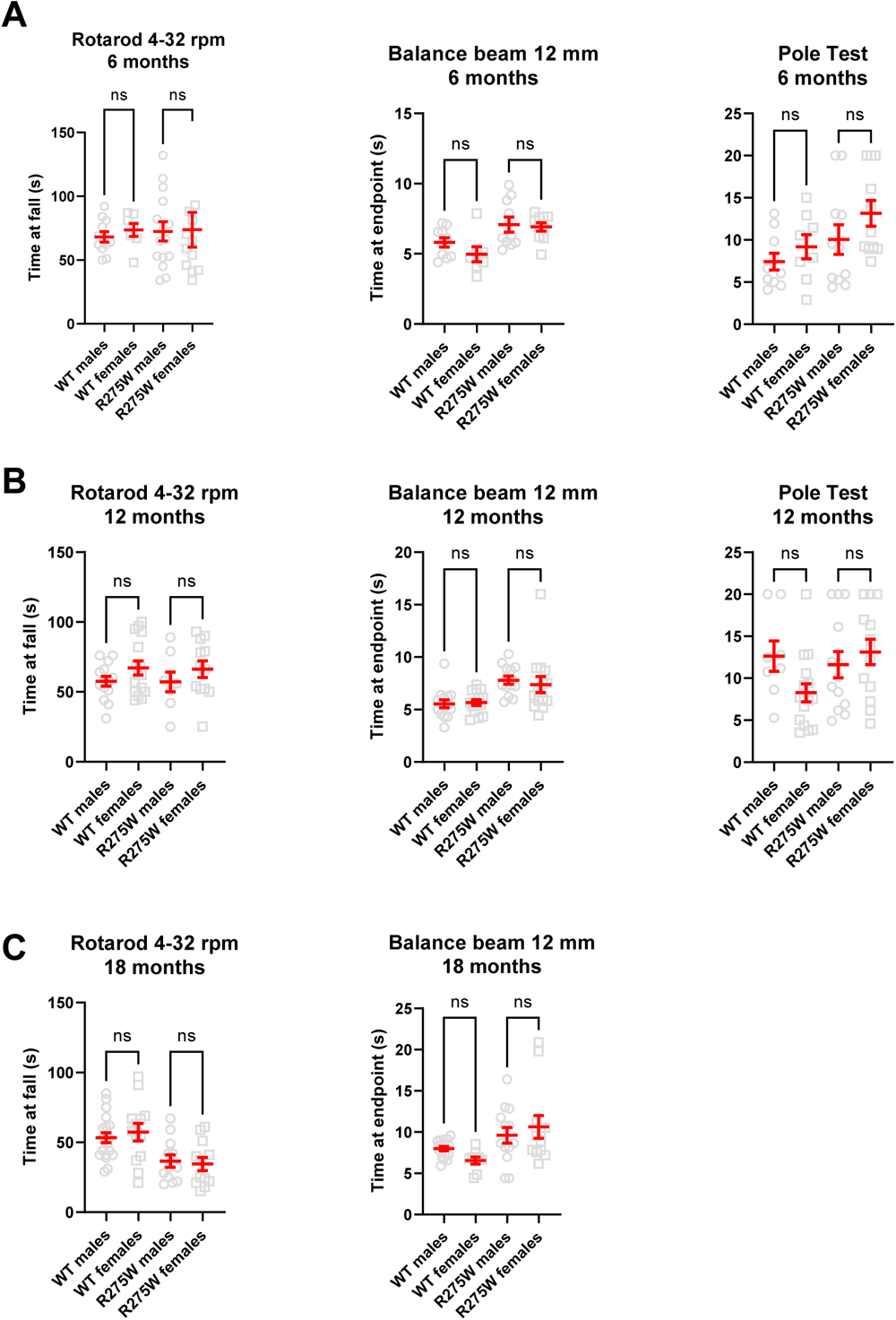
Data from motor tests analyzed in male and female. We tested WT and *Prkn*R275W mice at 6 months of age (A) 12 months of age (B) and 18 months of age (C).

