## supplemental files for "Dopamine neuron dysfunction and loss in the *Prkn*R275W mouse model of Juvenile Parkinsonism"

**Supplementary Materials**

**Caption:**

**1. Abbreviation for supplementary text (page 2)**

**2. Materials and methods (pages 3-11)**

**2. Supplementary Figures (page 12-19)**

**ABBREVIATIONS SUPPLEMENTARY TEXT**

β2M	beta-2 microglobulin

aCSF	artificial cerebrospinal fluid

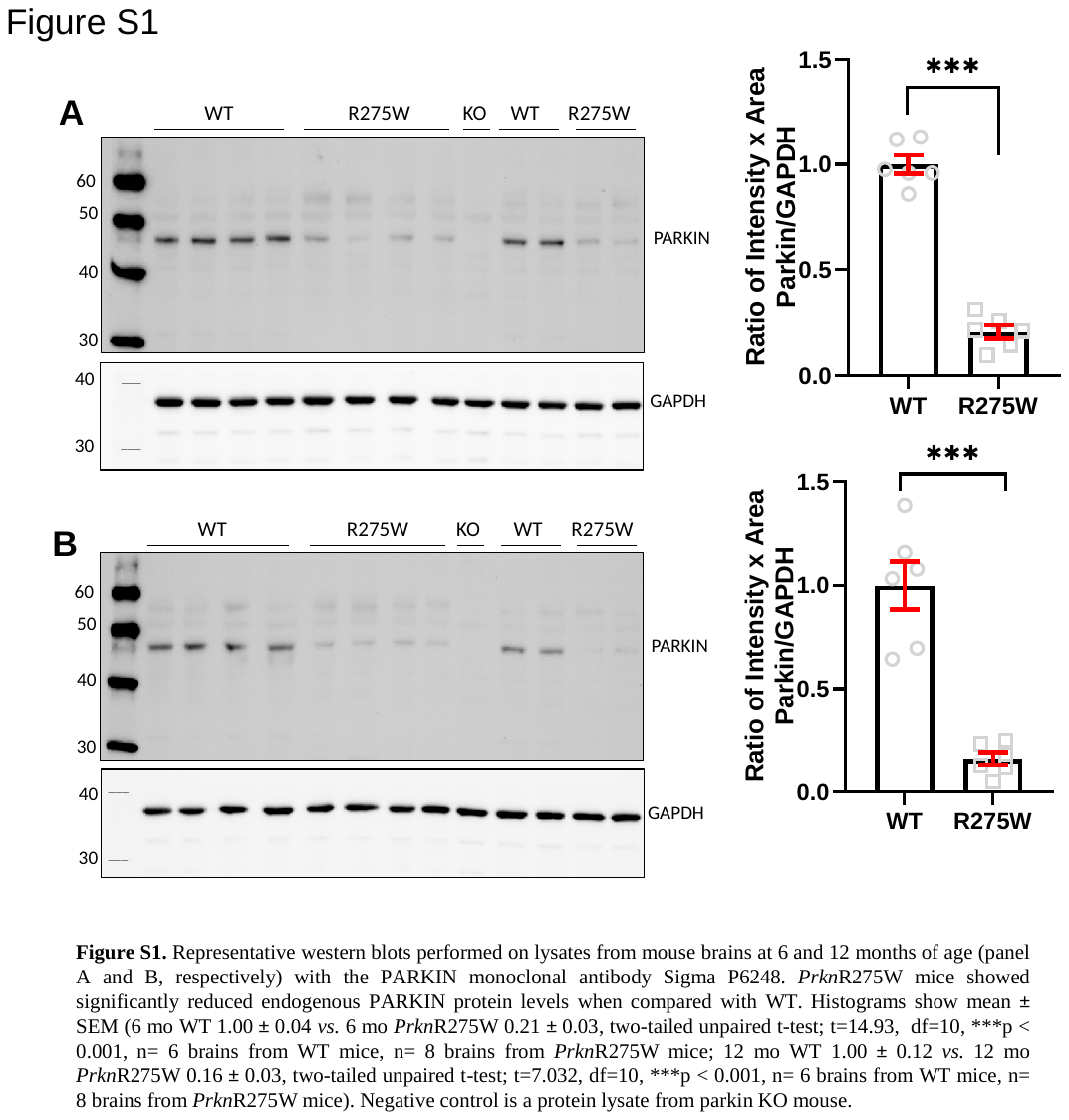

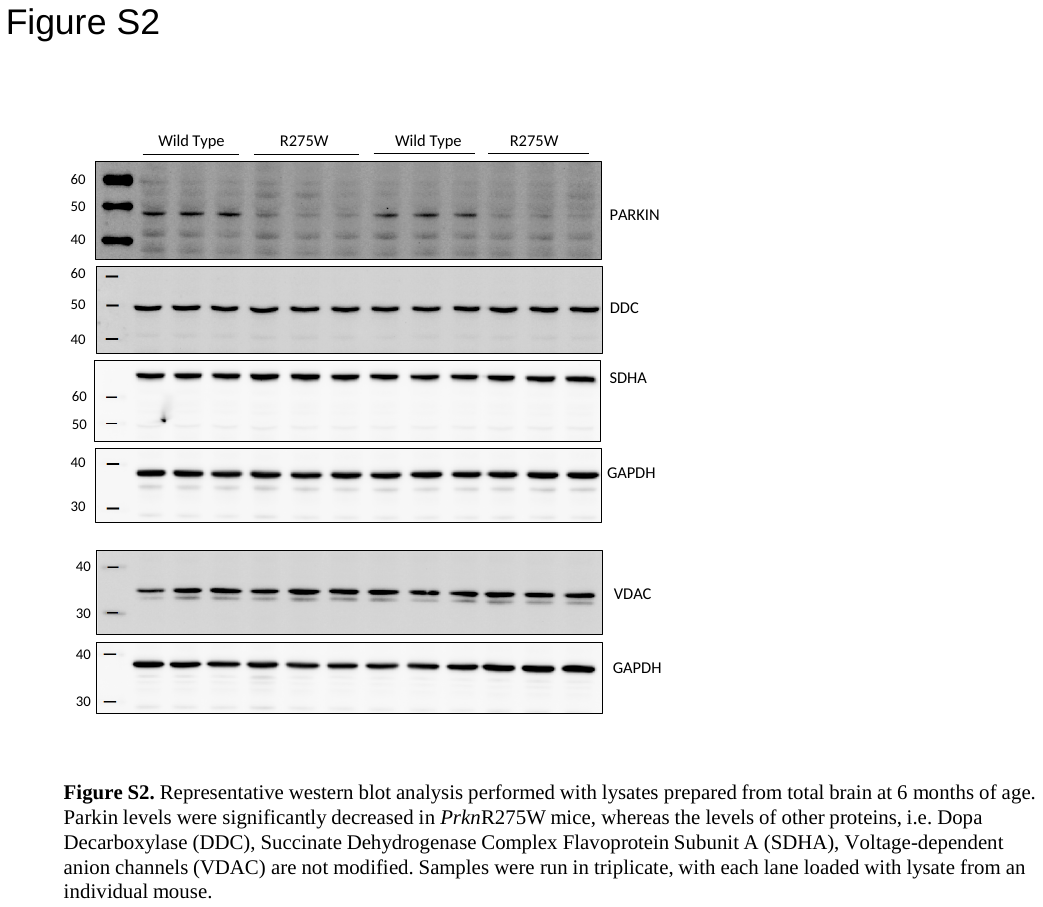

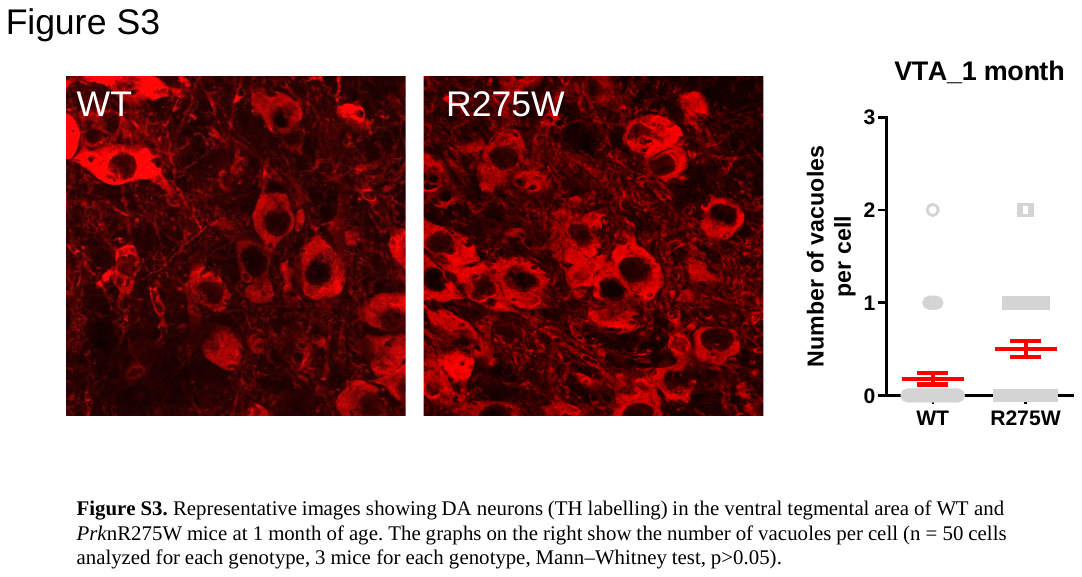

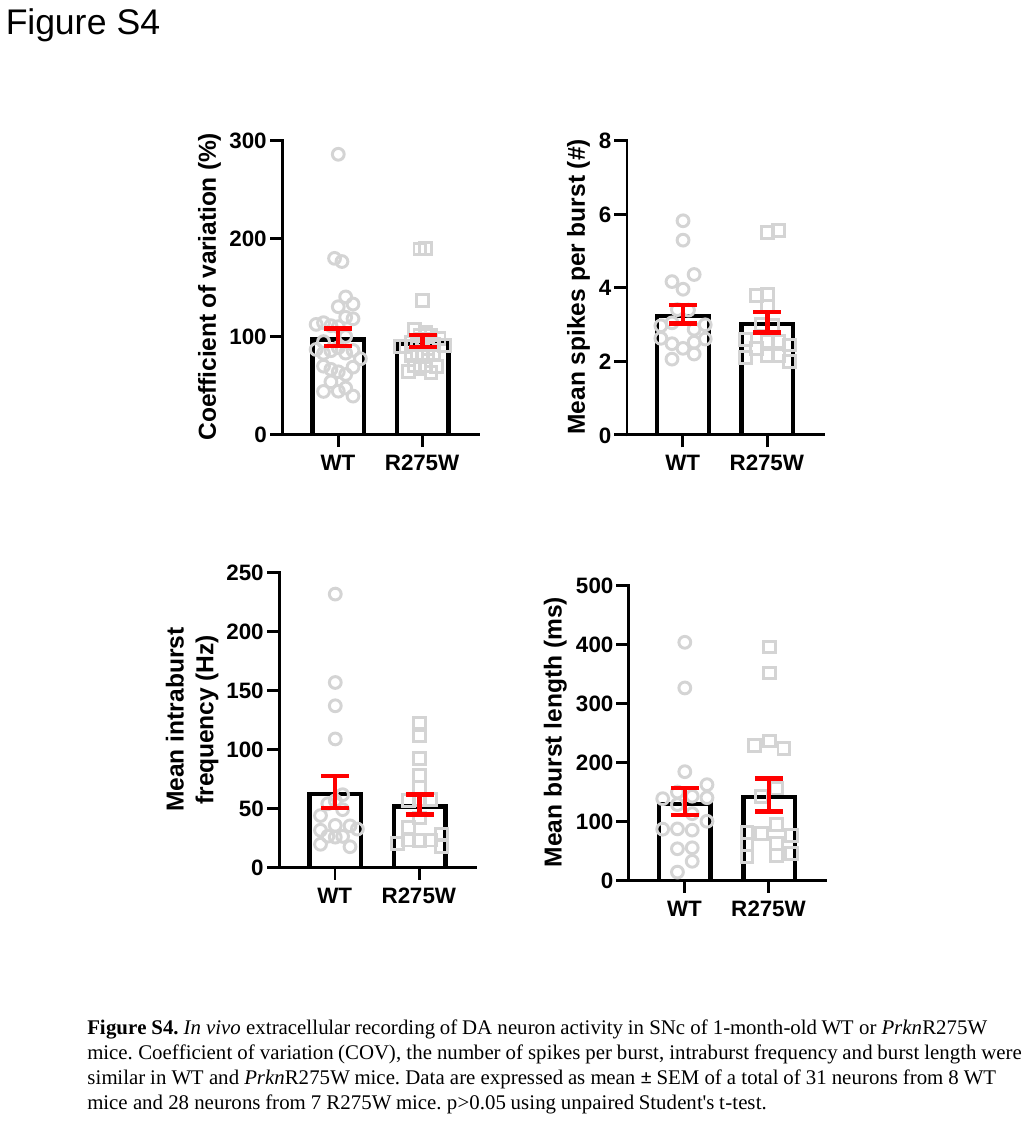

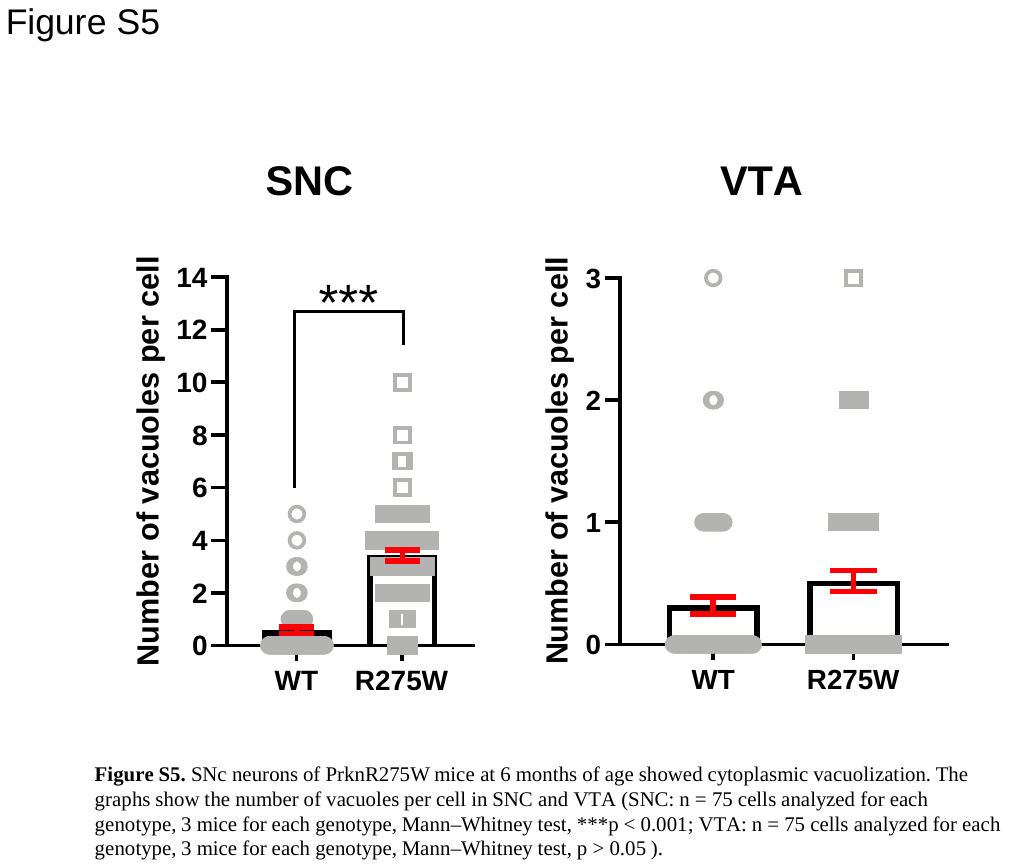

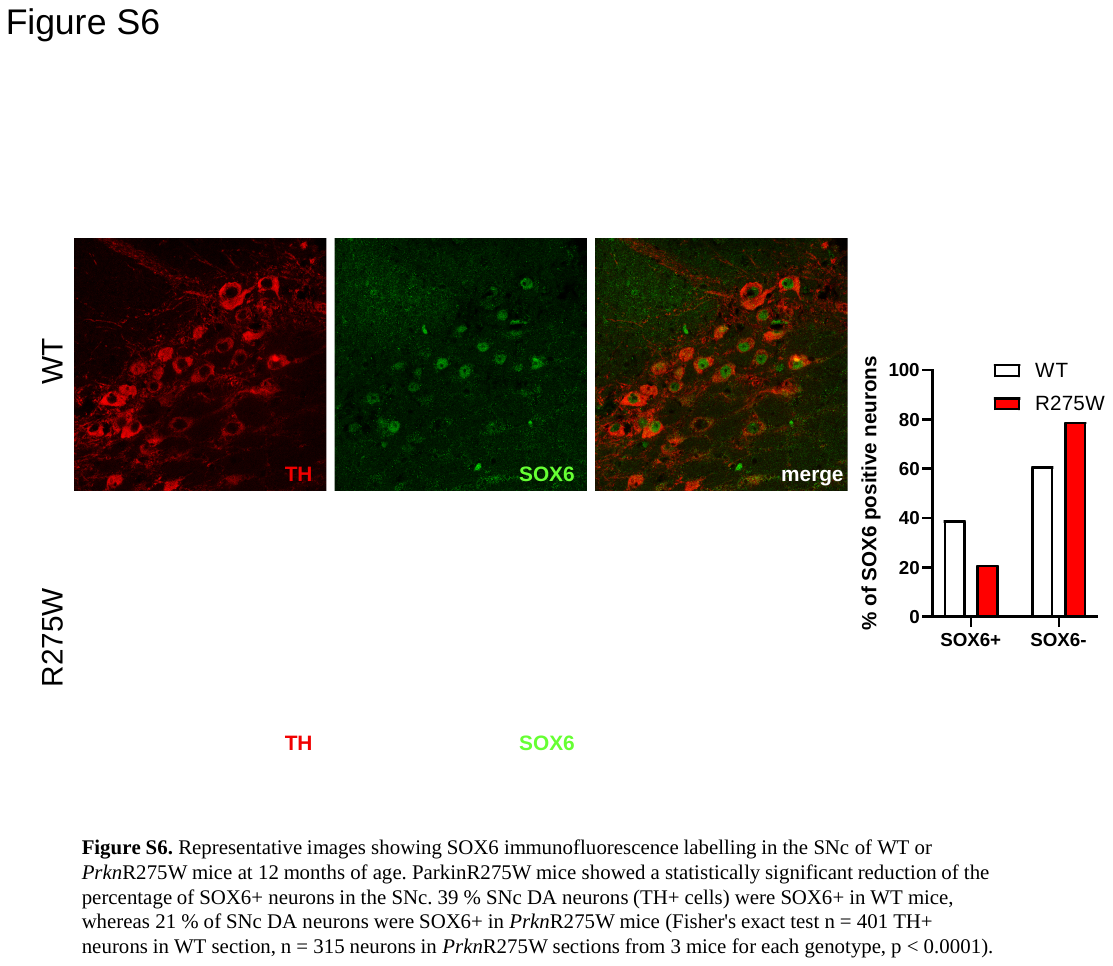

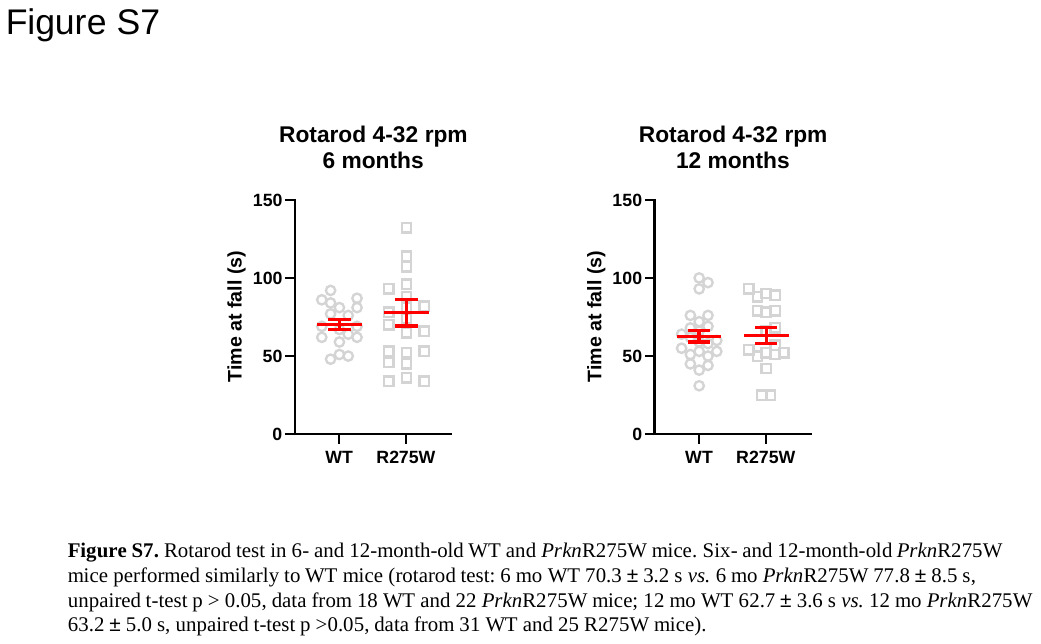

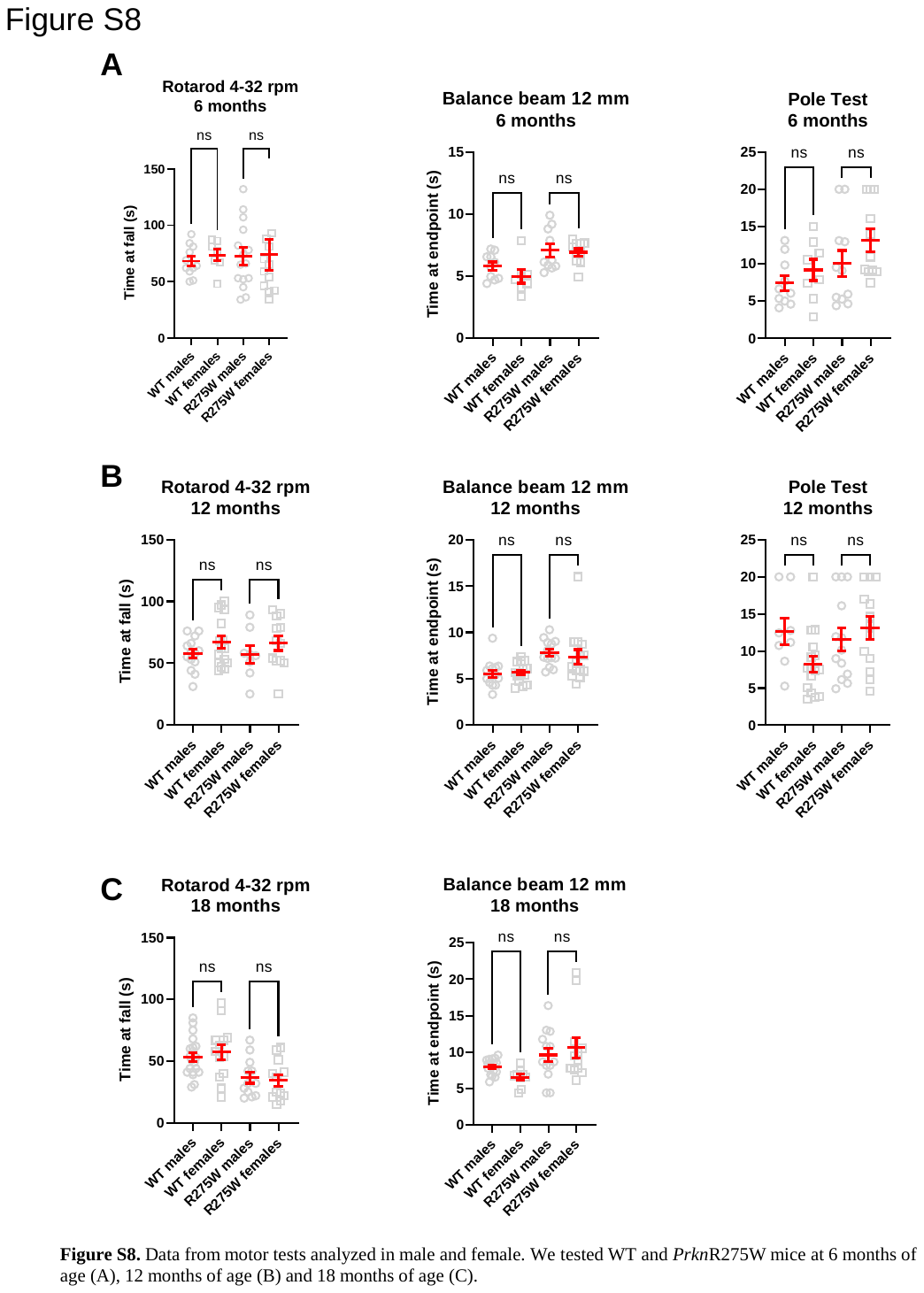
